# Autoinhibition of kinesin-1 is essential to the dendrite-specific localization of Golgi outposts

**DOI:** 10.1101/275149

**Authors:** Michael T. Kelliher, Yang Yue, Ashley Ng, Daichi Kamiyama, Bo Huang, Kristen J. Verhey, Jill Wildonger

## Abstract

Neuronal polarity relies on the selective localization of cargo to axons or dendrites. The molecular motor kinesin-1 moves cargo into axons but is also active in dendrites. This raises the question of how kinesin-1 activity is regulated to maintain the compartment-specific localization of cargo. Our in vivo structure-function analysis of endogenous Drosophila kinesin-1 reveals a novel role for autoinhibition in enabling the dendrite-specific localization of Golgi outposts. Mutations that disrupt kinesin-1 autoinhibition result in the axonal mislocalization of Golgi outposts. Autoinhibition also regulates kinesin-1 localization. Uninhibited kinesin-1 accumulates in axons and is depleted from dendrites, correlating with the change in outpost distribution and dendrite growth defects. Genetic interaction tests show that a balance of kinesin-1 inhibition and dynein activity is necessary to localize Golgi outposts to dendrites and keep them from entering axons. Our data indicate that kinesin-1 activity is precisely regulated by autoinhibition to achieve the selective localization of dendritic cargo.

**Summary:** Neuronal polarity relies on the axon-or dendrite-specific localization of cargo by molecular motors such as kinesin-1. These studies show autoinhibition regulates both kinesin-1 activity and localization to keep dendritic cargo from entering axons.

## Introduction

To receive and send signals, neurons rely on the polarized distribution of vesicles, macromolecules, and organelles to dendrites and axons. The majority of transport in neurons is mediated by the microtubule-based molecular motors kinesin and dynein. Dynein and kinesins traffic cargo in both compartments, raising the question of how motor function is regulated to maintain the polarized distribution of axonal and dendritic cargoes. To investigate this question, we focused on kinesin-1, which has been implicated in both axonal and dendritic transport.

Kinesin-driven events must be tightly controlled in the cell body to ensure that both outgoing and incoming cargos reach their proper destinations. In mammalian neurons, both axons and dendrites contain plus end-distal microtubules, indicating that kinesin-1 must be regulated to prevent the motor from taking cargo to the wrong compartment. One current model is that kinesin-1 is able to intrinsically distinguish between axonal and dendritic microtubules and that this recognition guides cargo-bound motors (Atherton et al., 2013; Bentley and Banker, 2016). This model arose from experiments characterizing the distribution of truncated, constitutively active kinesin motors lacking C-terminal cargo-binding and autoregulatory domains. The localization patterns of truncated kinesins in mammalian neurons and other cell types suggests that kinesin-1 and kinesin-3 differentiate between microtubule tracks patterned by post-translational modifications and microtubule-associated proteins (Atherton et al., 2013; Cai et al., 2009; Farias et al., 2015; Guardia et al., 2016; Hammond et al., 2010; Jacobson et al., 2006; Nakata and Hirokawa, 2003; Nakata et al., 2011; Tas et al., 2017; Tortosa et al., 2017). The adaptors that regulate motor-cargo attachment may also serve to regulate proper cargo distribution (Fu and Holzbaur, 2014).

In invertebrate neurons, sorting decisions in the cell body are theoretically simplified by the microtubule architecture as axons contain plus-end-distal microtubules whereas dendrites contain predominantly minus-end-distal microtubules (Rolls et al., 2007; Stone et al., 2008). However, for both vertebrate and invertebrate neurons, motor-driven transport must be highly regulated to prevent kinesin-driven retrograde transport in dendrites from continuing unimpeded into axons upon reaching the cell body. Similarly, in the cell body, kinesin-1 must be inactivated when bound to dendritic cargo before the cargo is transported out to dendrites.

Kinesin-1, like several other kinesin family members, is regulated by autoinhibition (Verhey and Hammond, 2009). Autoinhibition inactivates kinesin-1 when it is not cargo-bound and may fine-tune motor activity when kinesin-1 is cargo-bound (Fu and Holzbaur, 2013; Fu et al., 2014; Lane and Allan, 1999; Maday et al., 2014; Verhey and Hammond, 2009; Wozniak and Allan, 2006).

Although in vitro studies indicate that autoinhibition plays a key role in controlling motor function, the cellular role(s) for autoinhibition in regulating the function of kinesin-1, as well as other kinesins and dynein, are still poorly understood. To date, motor autoinhibition has been shown to modulate axonal transport (kinesin-1), axon outgrowth (kinesin-3), and synapse positioning (kinesin-4) (Cheng et al., 2014; Moua et al., 2011; Niwa et al., 2016; van der Vaart et al., 2013). Given the role of autoinhibition in these processes, it is attractive to consider the possibility that autoinhibition of kinesin-1 would regulate the compartment-specific localization of cargo as well.

We used in vivo structure-function analysis of endogenous, full-length kinesin-1 in fruit flies to uncover the mechanisms that regulate kinesin-1 activity to ensure the proper distribution of dendritic cargo. We sought to identify mutations that would lead to the axonal mislocalization of dendritic cargo, specifically Golgi outposts, which our data suggest are carried by kinesin-1. We initially tested the idea that kinesin-1 uses its microtubule-binding domain to “read” positional information encoded on microtubules. Studies of truncated kinesin-1 have implicated three different loops in the kinesin-1 microtubule-binding domain (β5-loop 8, loop 11, and loop 12) in polarized transport; however, there is no consensus on which loop(s) are essential (Huang and Banker, 2012; Konishi and Setou, 2009; Nakata and Hirokawa, 2003; Nakata et al., 2011). Our in vivo studies of full-length endogenous kinesin-1 show that E177 within β5-loop 8 is critical to the dendrite-specific localization of Golgi outposts, but does not affect the axon-specific distribution of presynaptic proteins including Bruchpilot, synapsin, and cysteine string protein.

This particular glutamate, E177, is part of a short motif previously implicated in kinesin-1’s preference for axons in vertebrate neurons (Huang and Banker, 2012; Konishi and Setou, 2009), but it also mediates contact with the C-terminal tail when kinesin-1 is in a folded autoinhibited conformation (Kaan et al., 2011). Our combined in vivo and in vitro results indicate that mutating E177 results in the axonal mislocalization of Golgi outposts by relieving kinesin-1 autoinhibition. We show that autoinhibition is also critical for the proper localization of kinesin-1 itself. Like constitutively active truncated motor constructs, uninhibited full-length kinesin-1 is enriched in axon terminals and nearly depleted from dendrites. The severe decease in kinesin-1 levels in dendrites correlates with the axonal mislocalization of Golgi outposts and a reduction in dendrite growth in the E177K mutant neurons. Golgi outposts are transported into dendrites by dynein (Lin et al., 2015; Ye et al., 2007; Zheng et al., 2008), and our genetic interaction tests suggest that a balance of dynein activity and kinesin-1 inhibition restricts outposts to dendrites and regulates their distribution within the arbor. Thus, our data indicate that the selective localization of cargo to dendrites relies on the autoinhibition of axon-targeted kinesin-1.

## Results

### Survival and neuronal morphogenesis depend on E177 in β5-loop 8 of kinesin-1

We initially set out to test whether full-length endogenous kinesin-1 is stopped from carrying dendritic cargo into axons via the recognition of microtubule-based signals. The highly conserved kinesin-1 microtubule-binding domain interfaces with microtubules via three loops. To identify the loop(s) essential to transport, we took an in vivo structure-function approach and mutated endogenous full-length kinesin-1 in fruit flies, which express a single highly conserved kinesin-1, Kinesin heavy chain (Khc). Most of what is known about kinesin-1-mediated transport is derived from analyzing truncated kinesin-1 constructs. Thus, it is unclear which domain(s) are necessary for the full-length motor to mediate the selective localization of cargo in vivo. We took advantage of a fly strain that we previously generated that enables the rapid knock-in of *Khc* alleles via site-directed integration (Fig. 1 A) (Winding et al., 2016). As *Khc* is an essential gene, we used animal viability as an initial screen to identify the loop(s) critical to Khc function.

**Figure 1:**
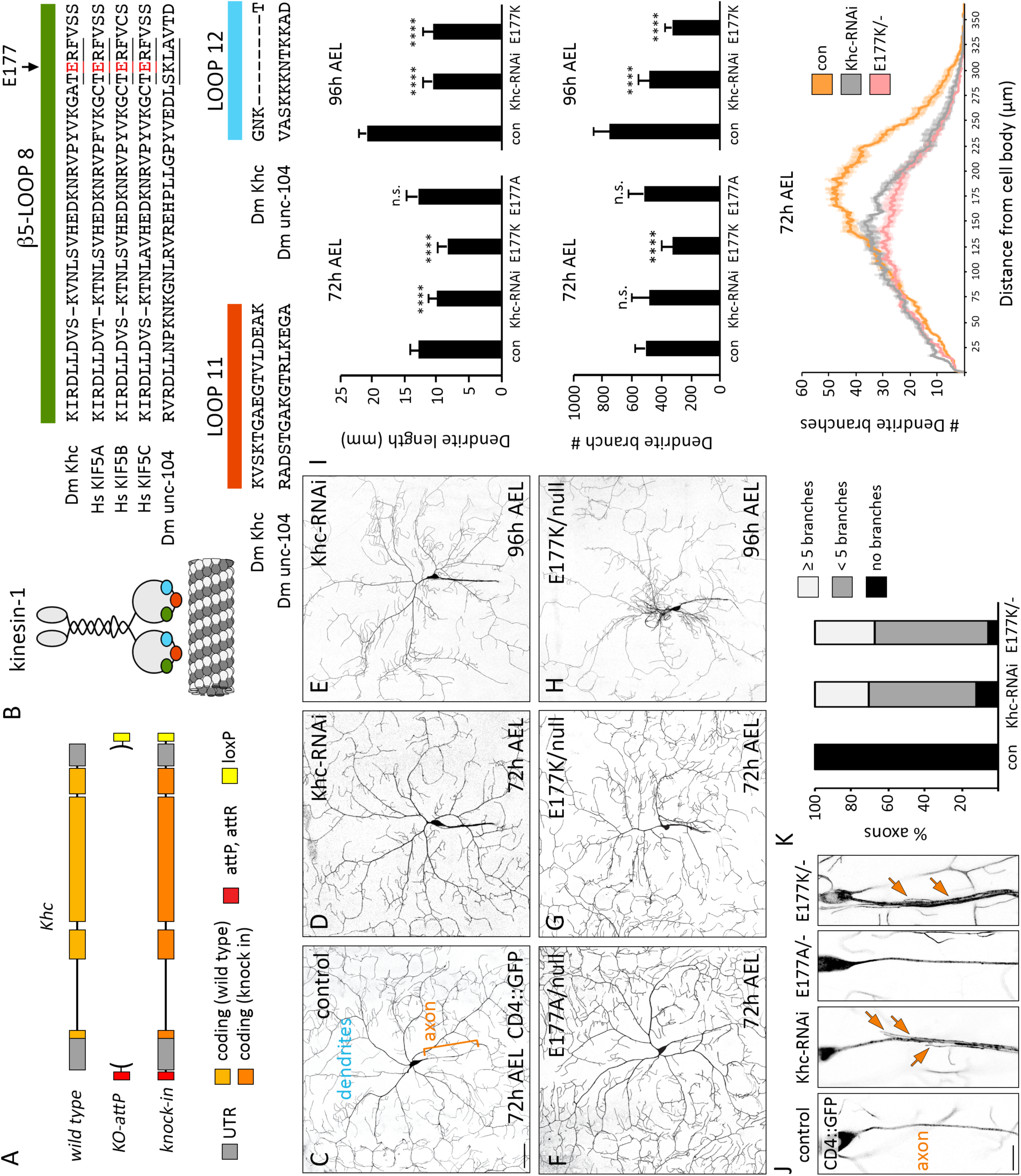
Dendrite arborization is disrupted by charge-reversal mutagenesis of E177 in 5-loop 8 of Khc. (A) Cartoon of the engineered *Khc* locus in which the entire Khc gene is replaced by an *attP* site to facilitate the rapid, reliable knock in of designer alleles with a minimal footprint (*attR, loxP* sites). (B) Cartoon of kinesin-1 dimer highlighting β5-loop 8 (green), loop 11 (orange), and loop 12 (blue) at the microtubule-binding interface. Alignment of Khc sequences that were substituted with the equivalent unc-104 sequences. Sequences of β5-loop 8 from three human kinesin-1 family members, KIF5A-C, are also shown to highlight the conservation between flies and humans. The TERF sequence is underlined; arrow indicates Khc E177, which is also highlighted in red. (C-I) Representative images (C-H) and quantification of dendrite length, branch number, and branch complexity (I) of control and mutant class IV dendritic arborization sensory neurons at 72 and 96 h AEL. Scale bar = 50 μm. Quantification of dendrite length and branch number (average ± SD) at 72 h AEL of 11 (*Khc* ^*E177A/-*^), 13 (*Khc* ^*E177K/-*^), and 16 (control and *Khc-RNAi*) neurons in at least three larvae and at 96 h AEL of 10 (control, *Khc-RNAi*, and *Khc* ^*E177K/-*^) neurons in at least three larvae. ^****^P < 0.0001 in comparison to control and evaluated by one-way ANOVA and Tukey post-hoc test. Sholl analysis of dendrite arbors at 72 h AEL reveals the number of branches at given distances from the cell body (average ± SEM). Critical radius and maximum branches reported in Table 2. (J) The axons of control and *Khc* ^*E177A/-*^ neurons are thin and unbranched whereas the axons of *Khc* ^*E177K/-*^ and *Khc-RNAi*-expressing neurons frequently sprout ectopic fine branches. Scale bar = 10 μm. (K) Number of axons with no branches, fewer than five branches, or five and more branches in control, *Khc-RNAi*, and *Khc* ^*E177K/-*^ neurons (n=18 axons for all genotypes).

We repeated the loop-swap experiments that previously pointed to β5-loop 8, loop 11, and loop 12 as being integral to polarized transport (Huang and Banker, 2012; Konishi and Setou, 2009; Nakata and Hirokawa, 2003; Nakata et al., 2011). We substituted these highly-conserved loops in Khc with the equivalent loops from kinesin-3 (Table 1, Fig. 1 B). Kinesin-3 is a structurally similar motor that is active in neurons, but, unlike kinesin-1, its truncated motor domain localizes to both axons and dendrites in mammalian neurons, suggesting it “reads” microtubule cues differently than kinesin-1 (Jacobson et al., 2006; Nakata and Hirokawa, 2003). In fruit flies, there are two kinesin-3 motors: unc-104 (also known as immaculate connections or imac) and Khc-73. We swapped the loops in Khc with the equivalent loops from unc-104, which has an essential role in neuronal transport (Barkus et al., 2008; Pack-Chung et al., 2007)(Fig. 1 B). Substituting loop 11 (*Khc* ^*loop-11-swap*^) did not affect animal survival, indicating that the unc-104 loop 11 is able to functionally substitute for Khc loop 11 (Table 1). *Khc* ^*loop-12-swap*^ animals were also homozygous viable; however, the *Khc* ^*loop-12-swap*^ mutation was lethal in trans to a null allele of *Khc*, suggesting that *Khc* ^*loop-12-swap*^ is hypomorphic (Table 1). In contrast to the loop-11 and loop-12 substitutions, swapping Khc Error! Not a valid embedded object. β5-loop 8 with unc-104 β5-loop 8 resulted in lethality (Fig.1 B, Table 1). This indicates that β5-loop 8 is essential to Khc function and that unc-104 β5-loop8 cannot functionally substitute for Khc β5-loop 8. β5-loop 8 is highly conserved between fly Khc and human KIF5A-C (Fig. 1 B). Four amino acids (TERF) in β5-loop 8 have been implicated in kinesin-1-mediated polarized transport and the glutamate in this motif contacts the tail when the motor is in a folded autoinhibited conformation, thus implicating this motif in both motor navigation and autoregulation (Kaan et al., 2011; Konishi and Setou, 2009). We substituted the TERF sequence in Khc with the equivalent sequences from unc-104 and Khc-73. Strikingly, substitution of Khc TERF with SKLA from unc-104 resulted in lethality, whereas substitution with the Khc-73 SQLA did not (Table 1). Notably, E177K had been previously uncovered in a genetic screen for lethal mutations in Khc (Brendza et al., 1999). We found that the charge-reversal mutations E177K and E177R caused lethality, whereas mutating E177 to alanine (E177A) had no effect (Table 1).

**Table 1:** Effect of kinesin-1 mutations on survival.

| Khc Allele | Description | Lethal Stage* | Notes |
| --- | --- | --- | --- |
| Control | Knock-in of wild type kinesin-1 sequence | Not lethal |  |
| Null | Q65-stop, protein null | 2<sup>nd</sup> instar | Also known as *Khc*<sup>27</sup> (Brendza et al., 1999) |
| Loop 12 Swap | Loop 12 swapped for unc-104 loop 12 | Not lethal | Lethal in trans to Khc null allele |
| Loop 11 Swap | Loop 11 swapped for unc-104 loop 11 | Not lethal | Not lethal in trans to Khc null allele |
| β5-loop 8 Swap | β5-loop 8 swapped for unc-104 β5-loop 8 | 2<sup>nd</sup> instar | Lethal in trans to Khc null allele |
| SKLA | TERF in loop 8 swapped for equivalent unc-104 sequence | 3<sup>rd</sup> instar |  |
| SQLA | TERF in loop 8 swapped for equivalent Khc-73 sequence | Not lethal |  |
| E177K | Charge reversal mutation of E177 in β5-loop 8 | 3<sup>rd</sup> instar | E177K knock-in recapitulates mutation in *Khc*<sup>19</sup> (Brendza et al., 1999) |
| E177R | Charge reversal mutation of E177 in β5-loop 8 | 3<sup>rd</sup> instar |  |
| E177A | Alanine substitution of E177 in β5-loop 8 | Not lethal | Not lethal in trans to Khc null allele |
| S246F | Mutation in loop 11 that disrupts ATP hydrolysis | Pupae | S246F knock-in recapitulates mutation in *Khc*<sup>17</sup> (Brendza et al., 1999) |
| E164K | Mutation in β5-loop 8 that disrupts ATP hydrolysis | 3<sup>rd</sup> instar | Also known as *Khc*<sup>23</sup> (Brendza et al., 1999) |
| Δhinge2 | Deletion of hinge 2 in stalk | 3<sup>rd</sup> instar | Disrupts autoinhibition, also known as KHCΔH2 (Barlan et al., 2013) |
| R947E | Charge-reversal mutation of R947 in the autoinhibitory IAK motif | 3<sup>rd</sup> instar | R947 and E177 form a salt bridge when the motor is autoinhibited (Kaan et al., 2011) |
| P945S | Mutation of P945S in the autoinhibitory IAK motif in the tail | 3<sup>rd</sup> instar | Also known as *Khc*<sup>22</sup> (Moua et al., 2011), P945S interacts with F179 when the motor is autoinhibited (Kaan et al., 2011) |
| K944E | Charge-reversal mutation of K944 in the autoinhibitory IAK motif | 3<sup>rd</sup> instar |  |
| E177K, S246F | Charge reversal mutation of E177 combined with S246 mutation that decreases ATP hydrolysis | Pupae |  |
| S181A, S182A | Alanine substitution of serines adjacent to TERF motif | Not lethal | Not lethal in trans to Khc null allele |
| S181D, S182D | Phosphomimetic mutation of serines adjacent to TERF motif | Not lethal | Lethal in trans to Khc null allele |

\*Stage of development when ≥ 75% of homozygous animals are dead.

**Table 2:**
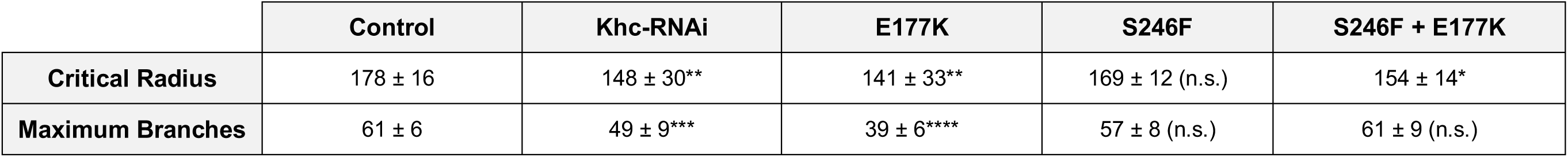
Effects of Khc knock-down and mutations on dendrite arborization. Sholl analysis of dendritic arbors quantifies the number of dendrites that intersect concentric circles centered on the cell body. The radius of the circle with the maximum dendritic intersections is the critical radius and the intersections at this radius are the maximum number of branches. Average ± SD for both measurements; n (neurons) = 14 control, 16 *Khc-RNAi*, 10 *Khc* ^*E177K*^, 14 *Khc* ^*S246F*^, 11 *Khc* ^*S246F,E177K*^. *P=0.05–0.01, ^**^P=0.01-0.001, ^****^P < 0.0001, n.s. = not significant in comparison to control and evaluated by one-way ANOVA with Tukey post-hoc test.

We next analyzed the effects of the Khc E177 mutations on neuronal morphogenesis. If the E177 mutations perturb dendritic transport, we expect to see a change in dendrite arborization. The Drosophila class IV dendritic arborization sensory neurons elaborate a highly branched dendritic arbor in the periphery and extend a single unbranched axon into the ventral nerve cord (VNC). Decreasing Khc levels reduced dendrite arborization and resulted in axons with minor ectopic branches (Fig. 1, C-E, and I-K), consistent with previous reports that Khc is essential for dendrite morphogenesis (Satoh et al., 2008). The viable *Khc* ^*E177A*^ mutation had no effect on dendrite or axon morphogenesis, but the *Khc* ^*E177K*^ mutants resembled *Khc-RNAi* neurons (Fig. 1, F-K and Table 2). The dendrite arborization phenotype of both the *Khc-RNAi* and *Khc* ^*E177K*^ neurons became more pronounced as the animals aged (Fig. 1 D, E, G, and H). These *Khc* ^*E177K*^ mutant phenotypes suggest that the E177K mutation disrupts kinesin-1’s function in neuronal morphogenesis.

### The Khc E177K mutation disrupts the dendrite-specific localization of Golgi outposts and hTfR-positive vesicles

The *Khc* ^*E177K*^ neurons resemble mutants in which dendritic transport is disrupted. Moreover, the ectopic axonal branching is characteristic of mutations that result in the axonal mislocalization of dendrite-specific Golgi outposts (Arthur et al., 2015; Satoh et al., 2008; Ye et al., 2007; Zheng et al., 2008). This led us to test whether Khc transports Golgi outposts and has a role in restricting them to dendrites. Kinesin-1 has been linked to Golgi and Golgi membranes via its adaptor kinesin light chain (Klc), which we found partially co-localizes with a subset of stationary and moving Golgi outposts in dendrites (Fig. 2, A-D) (Allan et al., 2002; Gyoeva et al., 2000; Marks et al., 1994; Wozniak and Allan, 2006). A third of motile Golgi outposts co-localize with Klc, and the majority of these Klc-positive Golgi outposts move retrograde at a velocity consistent with kinesin-mediated transport (Fig. 2 D). In control neurons, Golgi outposts localized to dendrites and were seldom found in axons (Fig. 2 B). Golgi outposts mislocalized to axons in *Khc* ^*E177K*,^ *Khc* ^*E177R*^, and *Khc-RNAi*-expressing neurons (Fig. 2, E and F). The mislocalization of outposts in *Khc* ^*E177K*^ neurons was rescued by the neuron-specific expression of wild-type Khc, indicating a cell autonomous requirement for Khc in outpost distribution. Mislocalized outposts were visible in both the proximal axon and axon terminals in the VNC (Fig. 2, G and H). Golgi outposts localized normally in the *Khc* ^*E177A*^ neurons. These data indicate Khc transports Golgi outposts in dendrites and that the E177K and E177R mutations disrupt motor function to cause their ectopic axonal localization.

**Figure 2:**
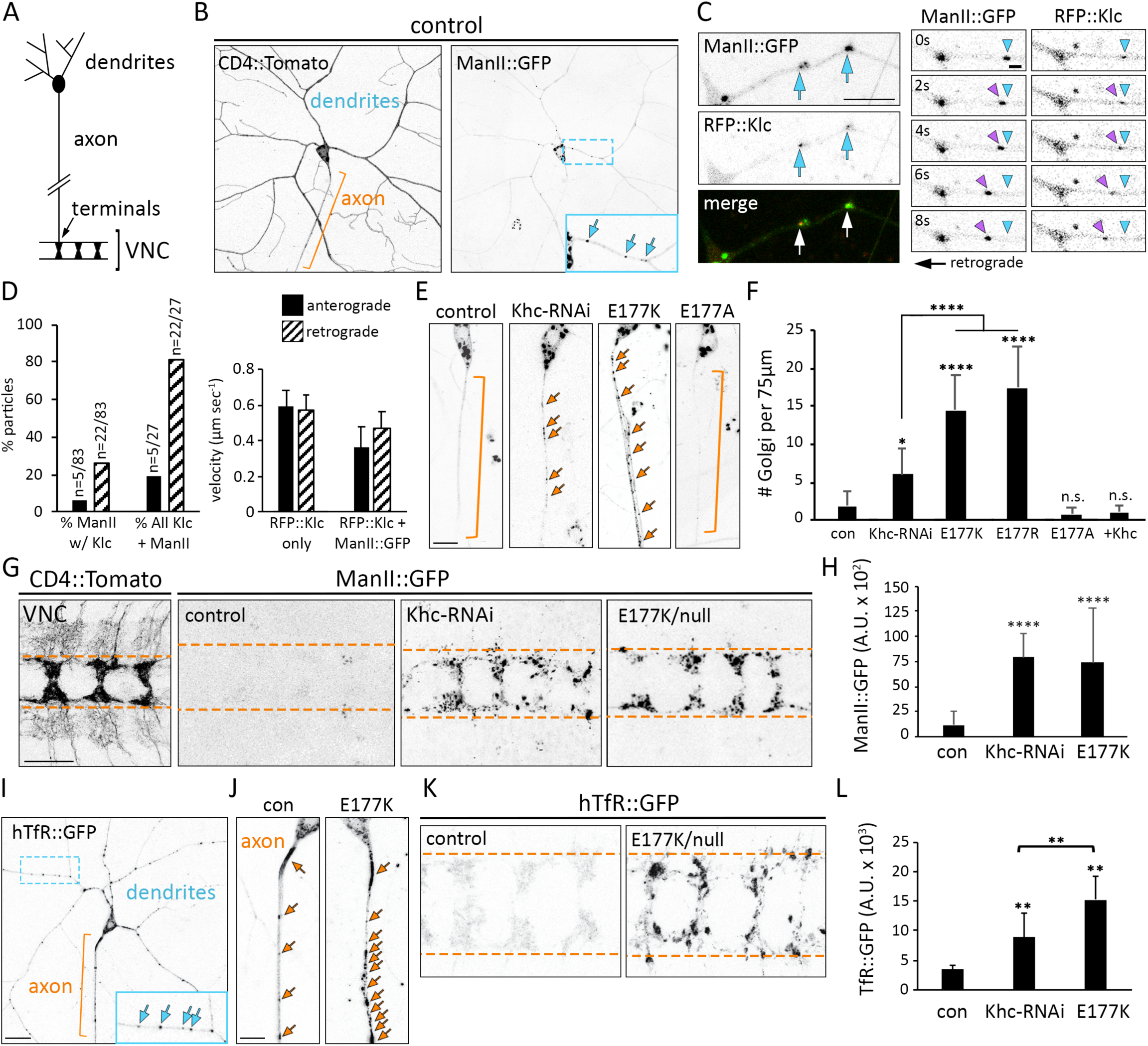
Ectopic axonal localization of dendritic Golgi outposts and hTfR-positive vesicles results from Khc E177 charge-reversal mutations. (A) Cartoon schematic of a class IV sensory neuron whose axon terminates in the VNC, where the class IV sensory neuron axons form a ladder-like pattern. (B) In control class IV sensory neurons, Golgi outposts labeled by ManII::GFP localize to dendrites and are excluded from axons. The zoomed inset (blue box) in the ManII::GFP panel shows several outposts in a dendrite branch. The membrane marker CD4::Tomato illuminates neuron morphology. Orange bracket indicates axon, blue arrows indicate Golgi outposts. (C) Co-localization of Golgi outposts and Klc in dendrites. Individual frames from a movie show a ManII-and Klc-positive punctum moving in a retrograde direction indicative of kinesin-mediated transport. Blue arrowhead indicates position at the start of the movie and purple arrowhead follows the punctum. Scale bar = 10 μm (left) and 2 μm (right). (D) Quantification of RFP::Klc-positive Golgi outposts. 32% of moving ManII::GFP-positive outposts (27/83) co-localize with RFP::Klc. Of these ManII::GFP-and RFP::Klc-positive outposts (n=27), 81% move retrograde at speeds consistent with kinesin-mediated transport. n=83 outposts and 75 RFP::Klc puncta in 24 neurons in 14 animals. The RFP::Klc only puncta may represent vesicles, which move faster than some organelles. (E) Representative images of Golgi outposts in control and mutant axons. Golgi outposts mislocalize to axons in *Khc* ^*E177K/-*^ and *Khc* ^*E177R/-*^ neurons, but not *Khc* ^*E177A/-*^ neurons (unbranched axons were selected for clarity). Expression of wild-type Khc (+Khc) in neurons rescues the mislocalization of outposts in *Khc* ^*E177K/-*^ larvae. Scale bar = 10 μm. (F) Quantification of Golgi outposts (average ± SD) in the proximal 75 μm of axons in 11 (*Khc* ^*E177A/-*^), 12 (*Khc* ^*E177K/-*^), 14 (*Khc* ^*E177R/-*^), and 15 (control, *Khc-RNAi*, and +Khc = *Khc* ^*E177K/-*^ *ppk-Gal4 UAS-Khc::BFP*) neurons in at least five larvae. (G) Images of class IV sensory neuron axons terminating in the VNC. CD4::Tomato reveals the normal ladder-like axonal projections. In control VNCs, the absence of ManII::GFP signal indicates Golgi outposts are normally excluded from axon terminals. Unlike control neurons, ManII::GFP accumulates in the axon terminals of *Khc* ^*E177K/-*^ and *Khc-RNAi* neurons. In the CD4::Tomato panel, the bushy signal flanking the axon terminals results from the non-specific expression of the transgene in a few central neurons. Scale bar = 25 μm. Dashed orange lines demarcate axon terminal borders in the VNC. (H) Quantification of ManII::GFP signal (average ± SD) for 25 (control, *Khc-RNAi*, and *Khc* ^*E177K/-*^) VNC segments in five larvae. (I) In control class IV sensory neurons, hTfR::GFP puncta localize to dendrites and are mostly excluded from axons. The zoomed inset in the hTfR::GFP panel shows several puncta in a dendrite branch. (J) Representative images of hTfR::GFP particles in control and mutant axons. hTfR::GFP mislocalizes to axons in the *Khc* ^*E177K/-*^ neurons (an unbranched axon was selected for clarity). Scale bar = 10 μm. (K) In control VNCs, the absence of hTfR::GFP signal indicates hTfR::GFP-containing vesicles are normally excluded from axon terminals. Unlike control neurons, hTfR::GFP accumulates at the axon terminals of *Khc* ^*E177K/-*^ and *Khc-RNAi* neurons. Arrows indicate hTfR::GFP puncta (I, J) and dashed lines demarcate axon terminal borders in the VNC (K). (L) Quantification of hTfR::GFP signal (average ± SD) in 25 (control, *Khc-RNAi*, and *Khc* ^*E177K/-*^) VNC segments in five larvae. ^*^P=0.05–0.01, ^**^P=0.01–0.001, ^****^P < 0.0001, and n.s. = not significant in comparison to control and evaluated by one-way ANOVA and Tukey post-hoc test (F, H, L).

To determine whether additional dendritic cargos would be affected by the Khc E177K mutation, we analyzed the localization of GFP-tagged human transferrin receptor (hTfR), a commonly used marker of dendritic vesicles that are transported, at least in part, by kinesin-1 and whose localization in fly neurons is Khc dependent (Henthorn et al., 2011; Schmidt et al., 2009). In control neurons, hTfR::GFP was enriched in dendrites with occasional puncta visible in axons and axon terminals (Fig. 2, I and J). The *Khc* ^*E177K*^ mutation resulted in an accumulation of hTfR::GFP in axons and axon terminals (Fig. 2, J-L). Although Golgi outposts and hTfR::GFP mislocalize to axons, we found no aberrant dendritic localization of three different axonal proteins (Fig. S1). Thus, Khc E177 has a critical role in restricting the localization of both Golgi outposts and hTfR::GFP-positive vesicles to dendrites and preventing their entry into axons.

### *Khc* ^*E177K*^ does not phenocopy mutations that disrupt kinesin-1 ATPase activity

The phenotypic similarity between *Khc* ^*E177K*^ and *Khc-RNAi* neurons initially suggested that the mislocalization of Golgi outposts in the E177K mutant may be due to decreased motor activity rather than a change in the ability of the motor to read out microtubule-based signals or in autoinhibition, which would likely increase motor activity. To test this possibility, we compared the effects of the E177K mutation to E164K and S246F mutations known to alter Khc ATPase activity (Brendza et al., 1999). The E177K mutation caused a dramatic mislocalization of Golgi outposts to axons yet had no effect on the motor’s velocity or microtubule dwell time in single-molecule motility assays (Fig. 3B and Fig. S2). In contrast, Golgi localization was unaffected by E164K or S246F mutations (Fig. 3, A and B) despite strong effects of these mutations on the motor’s landing rate and/or dwell time on microtubules (Fig. S2). Furthermore, the E177A mutation did not affect animal survival, dendrite morphogenesis, or Golgi outpost localization despite a demonstrated effect on microtubule-stimulated ATPase activity and velocity for a truncated mammalian motor domain (Woehlke et al., 1997). Thus, the E177K mutation is unlikely to disrupt Golgi outpost localization simply by diminishing Khc’s motility properties.

**Figure 3:**
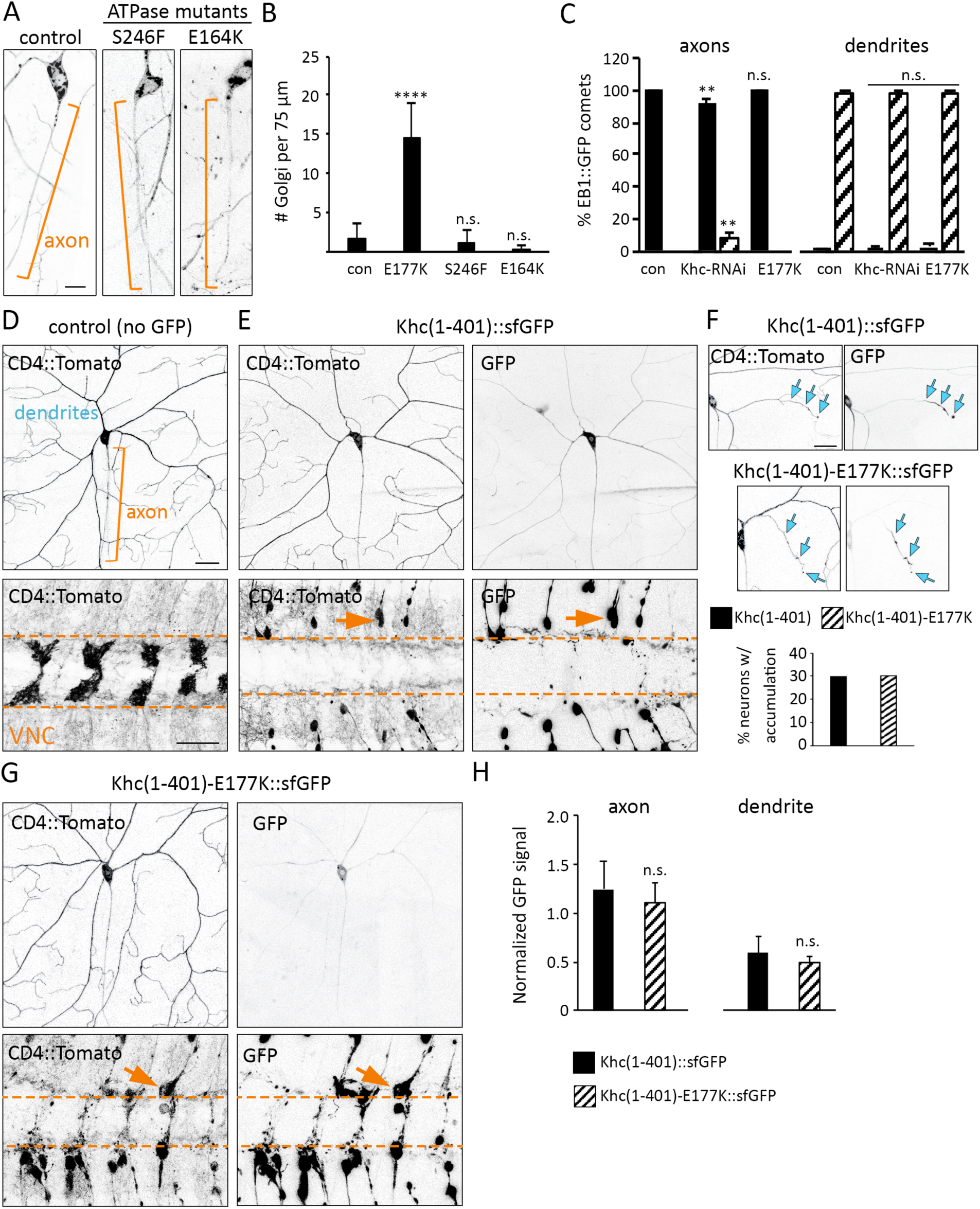
The Khc E177K mutation does not phenocopy ATPase mutants, disrupt microtubule polarity, or affect the localization of truncated constitutively active Khc. (A) Disruption of kinesin-1 ATPase activity by the S246F and E164K mutations does not cause the mislocalization of Golgi outposts to axons in contrast to the E177K mutation. Brackets indicate axons, arrows indicate Golgi outposts. Scale bar = 10 μm. (B) Quantification of Golgi outposts (average ± SD) in the proximal 75 μm of axons in 12 (*Khc* ^*E177K/-*^ and *Khc* ^*E164K/-*^), 13 (*Khc* ^*S246F/-*^), and 15 (control) neurons in at least four larvae. (C) Quantification of EB1::GFP comets in the axons (left) and dendrites (right) of class IV sensory neurons at 72 h AEL. Microtubule polarity, as indicated by the direction EB1::GFP comets travel, is unaffected by the Khc E177K mutation. The polarity of axonal microtubules in *Khc-RNAi*-expressing neurons is significantly altered. Comets in 10 (control and *Khc* ^*E177K/-*^) and 12 (*Khc-RNAi*) neurons were quantified. (D) CD4::Tomato reveals the morphology of the proximal dendrites and axon (top) as well as axon terminals in the VNC (bottom). (E) The truncated wild-type Khc construct, Khc(1-401)::sfGFP, is present in the cell body and localizes diffusely in the proximal axons and dendrites. Khc(1-401)::sfGFP strongly accumulates at the axon terminals. Khc(1-401)::sfGFP does not affect the morphology of the dendrites and axon, but causes axon terminals to retract (arrows). (F) Khc(1-401)::sfGFP and Khc(1-401)-E177K::sfGFP accumulate at one-to-three dendrite tips (arrows) in nearly a third of neurons. (G) Like the truncated wild-type motor, mutant Khc(1-401)-E177K::sfGFP is weakly visible in the proximal dendrites and axon and strongly accumulates in axon terminals, which retract similar to the Khc(1-401)::sfGFP. Scale bar = 25 μm (D-G). (H) Quantification of the GFP signal (normalized to the raw Tomato signal) in the proximal axons and dendrites for 22 [Khc(1-401)-E177K::sfGFP] and 30 [Khc(1-401)::sfGFP] neurons. ^**^P=0.01–0.001, ^****^P < 0.0001, n.s. = not significant in comparison to control and evaluated by one-way ANOVA and Tukey post-hoc test (B, C) or by a Student’s t-test (H).

### Axonal microtubule polarity is perturbed by *Khc-RNAi* but not the E177K mutation

The normal localization of Golgi outposts in the Khc ATPase mutants raises the question of why outposts mislocalize to axons when Khc is absent. Decreasing kinesin-1 levels or function can disrupt microtubule organization in axons and dendrites (Lee et al., 2017; Yan et al., 2013). In *Khc-RNAi*-expressing neurons the orientation of dendritic microtubules was normal, but the orientation of axonal microtubules was altered (Fig. 3 C). This change in axonal microtubule polarity may result in the ectopic axonal localization of outposts via dynein and/or another kinesin. The organization of microtubules in *Khc* ^*E177K*^ neurons was not affected, indicating that a change in microtubule polarity does not underlie the mislocalization of Golgi outposts in these neurons (Fig. 3 C). Ectopic axonal Golgi outposts have been speculated to disrupt the polarity of axonal microtubules due to their capacity to serve as platforms for microtubule nucleation (Ori-McKenney et al., 2012; Sanders and Kaverina, 2015). It is possible that the ectopic Golgi outposts in the *Khc* ^*E177K*^ axons may seed minus-end-distal microtubules, but that such microtubules are cleared by the mechanism(s) that normally maintain axonal microtubules in a uniform plus-end-distal array. Regardless, our data indicate that the ectopic outposts in *Khc* ^*E177K*^ axons are not a secondary consequence of a change in microtubule polarity.

### Localization of truncated constitutively active Khc is not affected by the E177K mutation

The effect of a mutation on a motor’s ability to navigate axonal and dendritic microtubules is typically tested using truncated motor constructs that lack cargo-binding and autoregulatory domains. We generated a truncated fly kinesin-1 construct, *Khc(1-401)::sfGFP*, that is equivalent to previously reported truncated vertebrate kinesin-1 constructs. Like truncated mammalian kinesin-1 (Nakata and Hirokawa, 2003), Khc(1-401)::sfGFP was present at low levels in the proximal axon and dendrites and accumulated in axon terminals (Fig. 3, D and E). Notably, Khc(1-401)::sfGFP accumulated in one or two dendrite branches in 30% of neurons (Fig. 3 F), presumably due to the presence of some plus-end-distal microtubules in developing dendrite branches. The axon terminals of *Khc(1-401)::sfGFP* neurons had retracted, indicating that persistent expression of truncated Khc(1-401) may have dominant-negative effect on axon growth (Seno et al., 2016). The mutant Khc(1-401)-E177K::sfGFP localized similarly to the wild-type truncated motor and there was no change in the percentage of neurons that accumulated Khc(401)-E177K::sfGFP in dendrites (Fig. 3, F-H). Thus, the E177K mutation does not noticeably disrupt the localization of the truncated, constitutively active motor.

### The E177K mutation disrupts autoinhibition of kinesin-1

Since our data suggest that the E177K mutation does not affect kinesin-1 function due to decreased ATPase activity or ability to navigate axonal and dendritic microtubules, we tested whether the E177K mutation might impinge on kinesin-1 autoinhibition. Khc E177 mediates an intramolecular salt bridge between the motor and tail domains when kinesin-1 is in its folded, autoinhibited state (Fig. 4 A) (Kaan et al., 2011). We reasoned that the E177K mutation disrupts the motor-to-tail interaction, preventing the motor from adopting the autoinhibited state.

**Figure 4:**
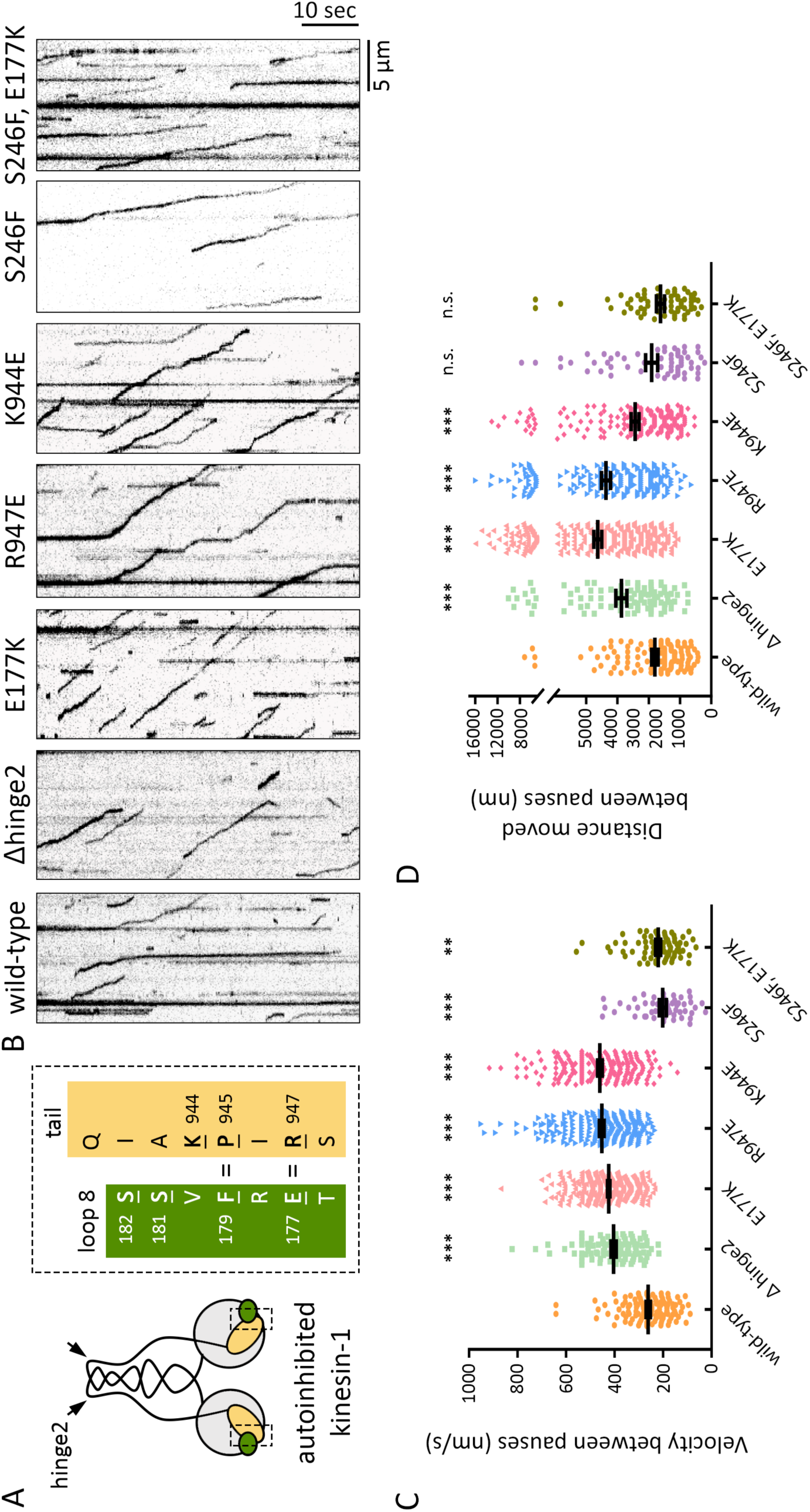
The E177K mutation increases Khc motility in single-molecule motility assays. Cartoon of a kinesin-1 dimer in the folded autoinhibited conformation in which β5-loop 8 in the motor domain (green) contacts the tail (yellow). Autoinhibition is mediated by interactions between residues in 5-loop 8 and the so-called IAK motif in the tail. Two critical intramolecular interactions occur between E177 and R947, and between F179 and P945. Residues that were mutated in our experiments are underlined. By eliminating a flexible hinge in the coiled-coil stalk, the Δhinge2 mutation prevents the motor from adopting the autoinhibited conformation. B)Representative kymographs showing single-molecule motility of full-length wild-type Khc and the indicated mutant versions. Microtubules are oriented plus-end to the right as in Fig. S2. All motors were tagged with mNeonGreen at their C-terminus and movies were acquired at 5 frames/sec. Scale bars = 5 μm (x-axis) and 10 sec (y-axis). (C,D) Quantification of velocities (C) and distances (D) between pauses (mean ± SEM) for each population of motors. Motility events analyzed: wild-type: 121, Δhinge2: 105, E177K: 366, R947E: 225, K944E: 209, S246F: 52 (velocity) and 60 (run length), and E177K,S246F: 79 (velocity) and 66 (run length). ^**^P=0.01-0.001, ^****^P < 0.0001, n.s. = not significant in comparison to the wild-type motor and evaluated by a two-tailed t-test.

To test this idea, we first turned to an in vitro single-molecule approach. Wild-type or mutant full-length Khc motors tagged with mNeonGreen were added to flow cells containing polymerized microtubules and their motility was observed by total internal reflection fluorescence (TIRF) microscopy. As expected, very few motility events were observed for wild-type Khc, which is typically in an autoinhibited state in solution (Fig. 4 B). When motile events were observed, the motor typically moved only a short distance at a slow speed before pausing or detaching (Fig. 4, B-D). As a positive control, we deleted the central hinge (Δhinge2) that enables the motor to fold on itself and adopt the autoinhibited state (Friedman and Vale, 1999). The Δhinge2 motors showed smooth movement along the microtubules, with few pauses, and with speeds typical of an active kinesin-1 motor (Fig. 4, B-D), consistent with a previous report (Friedman and Vale, 1999). Charge-reversal mutations to residues involved in maintaining the autoinhibited state (E177K, R947E, and K944E) (Hackney and Stock, 2000) also showed increased motility as compared to the wild-type motor with speeds characteristic of an active kinesin-1 motor and an increased distance moved along the microtubule track (Fig. 4. B-D). The mutant motors displayed more pauses during motility as compared to the Δhinge2 motor, likely because the motors with single mutations can still adopt the folded conformation whereas the Δhinge2 exists in the extended state. If the E177K mutation is relieving autoinhibition, then the resultant activation should be ameliorated by decreasing ATPase activity. We tested this idea by combining the E177K and S246F mutations. By itself the S246F mutation decreased motor speed without affecting the distance moved (Fig. 4, B-D). The double-mutant E177K, S246F motor behaved similarly to the single-mutant S246F and wild-type motors, indicating that reducing motor activity suppresses the effects of the E177K mutation. These findings support the idea that E177 has a role in regulating kinesin-1 autoinhibition and motor activation to control motor function.

### Disrupting autoinhibition perturbs Golgi outpost localization and dendrite morphogenesis

Given the single-molecule evidence that E177K disrupts kinesin-1 autoinhibition, we next asked whether other kinesin-1 mutations known to alter autoinhibition perturb the dendrite-specific localization of Golgi outposts. Indeed, similar to the Khc E177K mutant, neurons expressing the mutant Δhinge2 motor had Golgi outposts in axons (Fig. 5, A and B). Likewise, mutation of residues in the highly conserved autoinhibitory motif in the tail (so-called IAK motif) also disrupted outpost localization (Fig. 5, A and B). A charge reversal mutation of Khc R947, the tail residue that forms a salt bridge with E177, caused Golgi outposts to mislocalize to axons, as did additional mutations targeting other IAK motif residues (Fig. 5, A and B). Like the E177K mutation, the hinge2 and K944E mutations significantly reduced dendrite arborization (Fig. 5, C-G).

**Figure 5:**
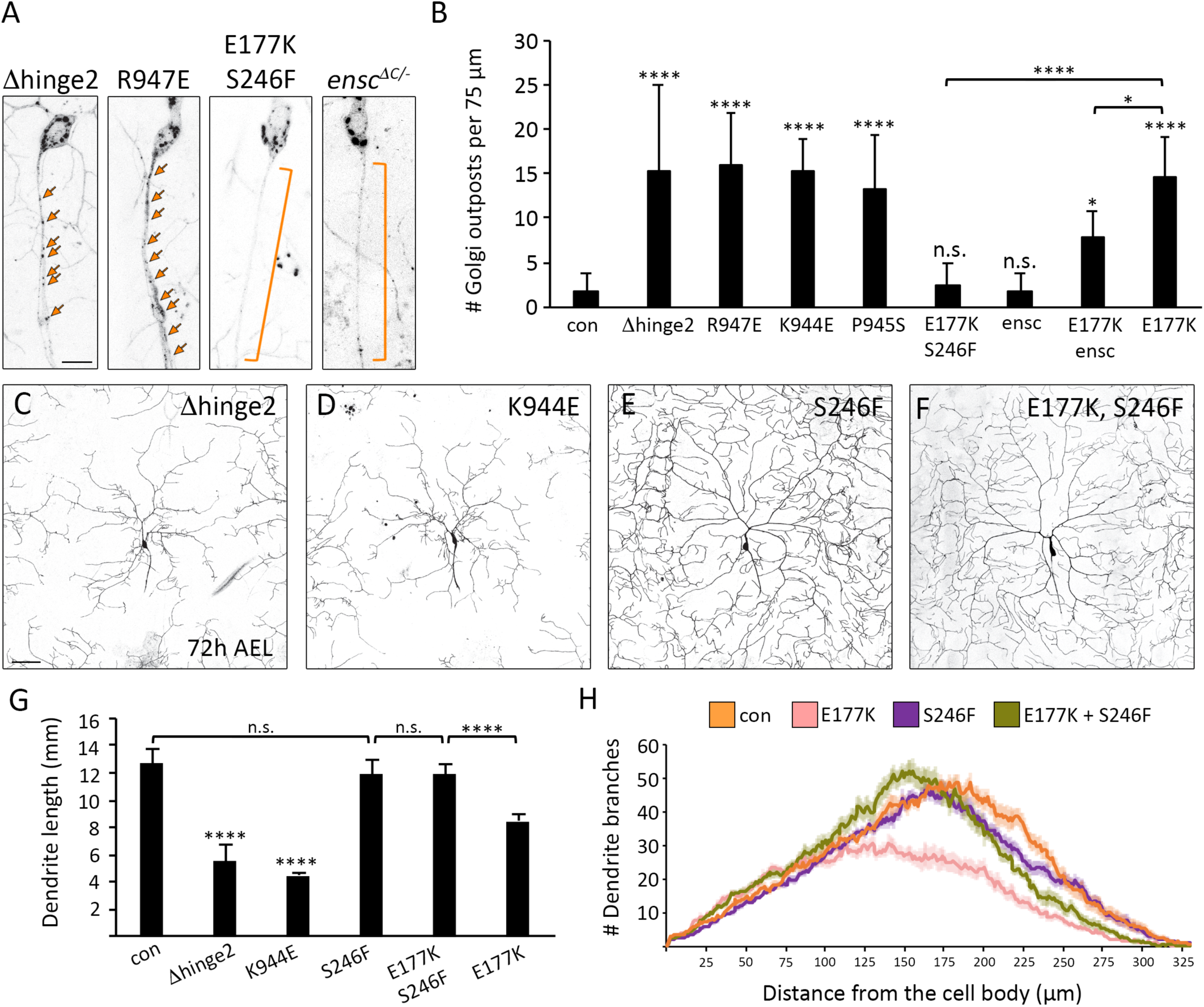
Khc mutations that disrupt autoinhibition result in the axonal mislocalization of Golgi outposts and perturb dendrite arborization. (A) Mutations that disrupt autoinhibition (Δhinge2, K944E, R947E, and P945S) result in the mislocalization of Golgi outposts to axons. The S246F mutation rescues the ectopic localization of Golgi outposts caused by the E177K mutation. The loss of ensconsin (*ensc* ^*ΔC/-*^), a kinesin-1 co-factor that relieves autoinhibition, does not affect the dendrite-specific localization of outposts and partially rescues the ectopic localization of Golgi outposts caused by the E177K mutation. Bracket indicates axon and arrows indicate Golgi outposts. Scale bar = 10 μm. (B) Quantification of Golgi outposts (average ± SD) in the proximal 75 μm of axons in 12 (*Khc* ^*Δhinge2/-*^ and *Khc* ^*E177K/-*^),13 (*Khc* ^*R947E/-*^ and *Khc* ^*P945S/-*^), 14 (*Khc* ^*K944E/-*^ and *ensc* ^*ΔC/-*^) and 15 (control, *Khc E*^177*K*,S246*F/-*^, and *Khc* ^*E177K/-*^, *ensc* ^*ΔC/-*^) neurons in at least five larvae. (C-H) Representative images (C-F) and quantification (G, H) of dendrite arbors in control and mutant neurons at 72 h AEL. Scale bar = 50 μm. Mutations that disrupt autoinhibition (Δhinge2, K944E) result in smaller dendritic arbors (C, D, G). The S246F mutation does not affect dendrite morphology (E, G, H). In *Khc* ^*E*177*K,S*246*F/-*^ double-mutant neurons, the S246F mutation rescues the decrease in arborization caused by the E177K mutation (F-H).Quantification (G) of total dendrite length (average ± SD) and Sholl analysis (H) of branch distribution (average ± SEM) in 13 (*Khc* ^*E177K,S246F/-*^ and *Khc* ^*E177K/-*^), 15 (*Khc* ^*Δhinge2/-*^, *Khc* ^*K944E/-*^, and *Khc* ^*S246F/-*^) and 16 (control) neurons in at least four larvae (G). The critical radius and maximum branches determined by Sholl analysis are reported in Table 2. *P=0.05–0.01, ^****^P < 0.0001, n.s. = not significant in comparison to control and evaluated by one-way ANOVA and Tukey post-hoc test (G, H).

These results suggest that ectopic kinesin-1 motor activity may be responsible for the change in Golgi outpost distribution. Our single-molecule results show that the S246F mutation, which slows ATPase activity, suppresses the increase in motor activity caused by the E177K mutation. We tested whether the S246F mutation would similarly suppress the mislocalization of Golgi outposts caused by the E177K mutation, and so we knocked-in a double-mutant E177K, S246F mutant motor. The *Khc* ^*E177K,S246F*^ neurons did not have any mislocalized Golgi outposts in their axons and had relatively normal dendrite arbors, indicating that an increase in Khc activity underlies the mislocalization of Golgi outposts and dendrite growth defect. (Fig. 5, A, B, and E-H and Table 2). We further tested this idea by decreasing the levels of the kinesin-1 co-factor ensconsin in the E177K mutant neurons. Ensconsin (also known as MAP7) relieves autoinhibition in Drosophila neurons and embryos to promote kinesin-1 activation (Barlan et al., 2013; Sung et al., 2008). Loss of ensconsin by itself did not disrupt Golgi outpost localization, but loss of ensconsin partially suppressed the mislocalization of Golgi outposts in the *Khc* ^*E177K*^ neurons (Fig. 5, A and B). These data support the idea that a less inhibited, more active kinesin-1 ectopically transports Golgi outposts into axons.

We also tested whether Golgi outpost distribution was affected by mutating two conserved serines near E177 (S181 and S182) whose phosphorylation has been implicated in regulating kinesin-1 autoinhibition (Morfini et al., 2009). Golgi outpost localization and dendrite arborization were altered by phosphomimetic mutations but not alanine mutations (Fig. S3). This suggests that phosphorylation of these serine(s) may favor the uninhibited active state whereas dephosphorylation may be permissive and allow the motor to switch between active and inhibited states.

### Loss of autoinhibition enriches Khc in axon terminals and reduces levels in dendrites

Since the E177K mutation makes kinesin-1 more active by relieving autoinhibition, we reasoned that the mutant full-length motor might localize primarily to axons similar to the truncated constitutively active motor. To test this idea, we tagged the endogenous wild-type and mutant Khc motors with the split-GFP peptide sfGFP11 (Cabantous et al., 2005; Kamiyama et al., 2016). We used split-GFP because the broad expression of *Khc* makes it difficult to assess the sub-cellular localization of GFP-tagged Khc in individual neurons. By restricting the expression of GFP1-10 to the class IV sensory neurons, we can readily visualize the reconstituted sfGFP in individual neurons and thus analyze the sub-cellular distribution of endogenous Khc. In the class IV sensory neurons, endogenous full-length Khc localizes to both dendrites, axons, and axon terminals (Fig. 6, A and E). The E177K mutation results in a notable and significant shift in Khc distribution from dendrites to axon terminals (Fig. 6, B, D-F). If this change in Khc localization is due to increased activity, the S246F mutation should restore its distribution.

**Figure 6:**
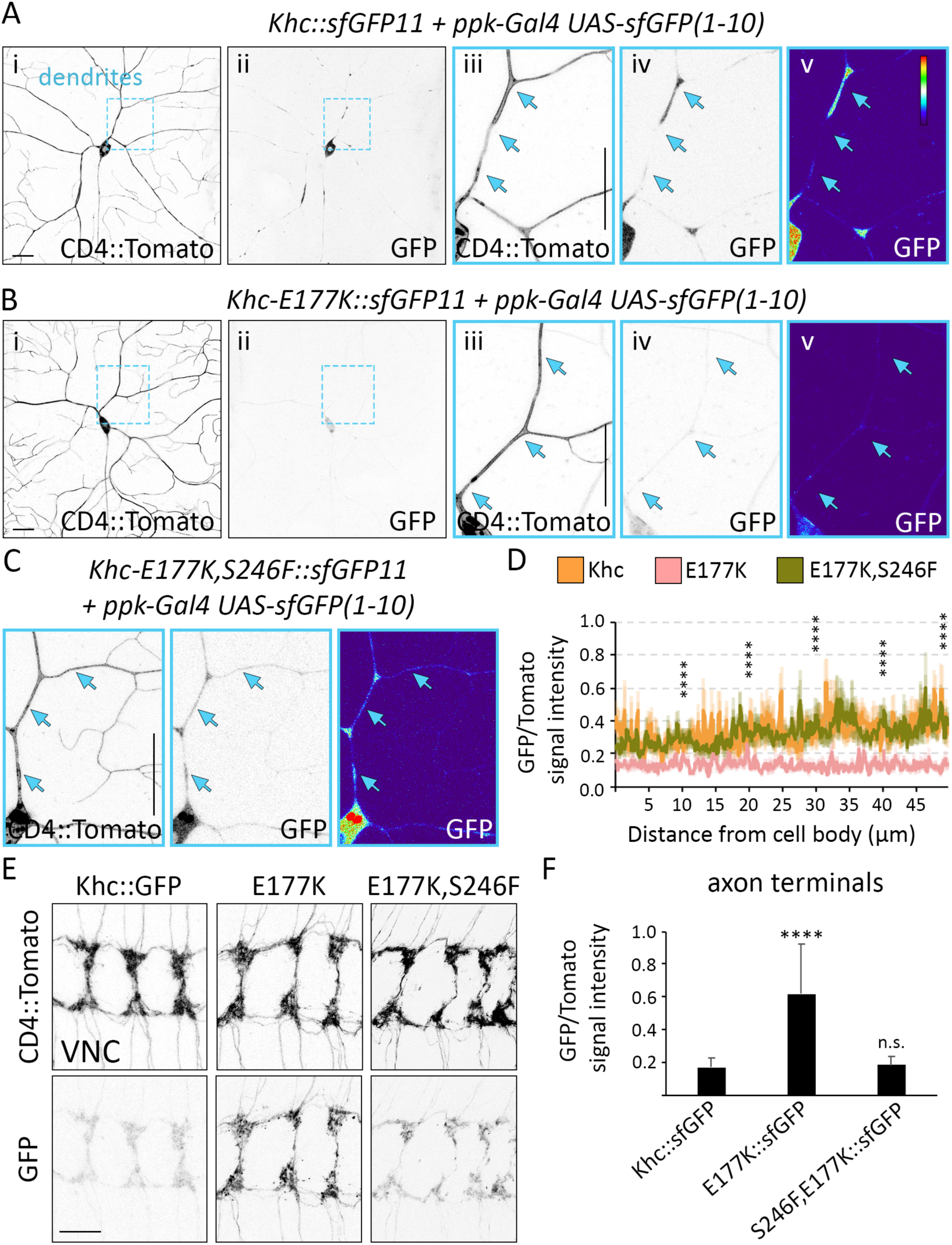
Disrupting Khc autoinhibition results in axonal accumulation of the motor. A split-GFP approach was used to visualize endogenous Khc. Khc was tagged with sfGFP11 and pickpocket (ppk)-Gal4 drove the expression of UAS-sfGFP(1-10) resulting in the class IV sensory neuron-specific reconstitution of sfGFP. (A-C) In contrast to the localization pattern of wild-type Khc::sfGFP (A), Khc-E177K::sfGFP (B) is significantly reduced in cell bodies and dendrites. Introducing the S246F mutation (Khc-E177K,S246F::sfGFP) restores the localization pattern to normal (C). (D) GFP signal intensity normalized to CD4::Tomato (average ± SEM) along the proximal 50 μm of the dendrite with the brightest GFP signal for 11 (Khc::sfGFP) and 12 (Khc-E177K::sfGFP and Khc-E177K,S246F::sfGFP) neurons in at least four larvae. The GFP/Tomato intensity ratios were compared at 10-μm intervals. Khc-E177K::sfGFP was significantly lower than Khc::sfGFP, but Khc-E177K, S246F::sfGFP was not statistically distinct from Khc::sfGFP at any interval. (E, F) Representative images of sfGFP-tagged wild-type and mutant motors reveal that Khc-E177K::sfGFP accumulates at axon terminals in the VNC (E). The S246F mutation rescues this abnormal accumulation of motor at terminals. Quantification of the GFP/Tomato intensity ratios at axon terminals (average ± SD) for 25 (Khc-E177K::sfGFP) and 29 (Khc::sfGFP and Khc-E177K,S246F::sfGFP) VNC segments in at least five larvae. ****P< 0.0001 and n.s. = not significant when compared to Khc::sfGFP and evaluated by one-way ANOVA and Tukey post-hoc test (D, F).

Indeed, the double-mutant E177K, S246F motor localized normally in both dendrites and axons (Fig. 6, C-F). This supports the idea the E177K mutation activates full-length Khc, virtually depleting the motor from dendrites and resulting in its axonal enrichment. The axonal shift in Khc distribution caused by the E177K mutation is consistent with the idea that the less inhibited mutant motor actively pulls Golgi outposts into axons.

### Reduced motility of Golgi outposts and hTfR-positive vesicles in E177K mutant dendrites

A significant decrease in dendritic Khc levels caused by the E177K mutation is likely to disrupt outpost motility. In support of this idea, mutations that disrupt autoinhibition reduce the fraction of motile outposts and increase the pause duration of the moving fraction. (Fig. 7, A-C). The E177K mutation significantly reduced the frequency of retrograde events and the likelihood that outposts moving anterograde switched direction to travel retrograde, consistent with a reduction of Khc in dendrites (Fig. 7 C). A significant decrease in anterograde reversals likely underlies the increased anterograde event frequency in the *Khc* ^*E177K*^ neurons (Fig. 7 C). Although pause duration increased, run length was normal in the mutants (Fig. S4 A). Golgi outposts were not the only cargo affected: the E177K mutation and expression of *Khc-RNAi* also decreased hTfR::GFP flux, most strikingly retrograde flux (Fig. 7 D and Fig. S4 B). In contrast to the autoinhibition mutants and knocking-down Khc, the ATPase mutant S246F had little effect on Golgi outpost or hTfR::GFP motility (Fig. 7, B-D). This suggests that the Khc S246F mutant is sufficiently active to support transport in dendrites, which likely explains why dendrite growth is not affected by this mutation.

**Figure 7:**
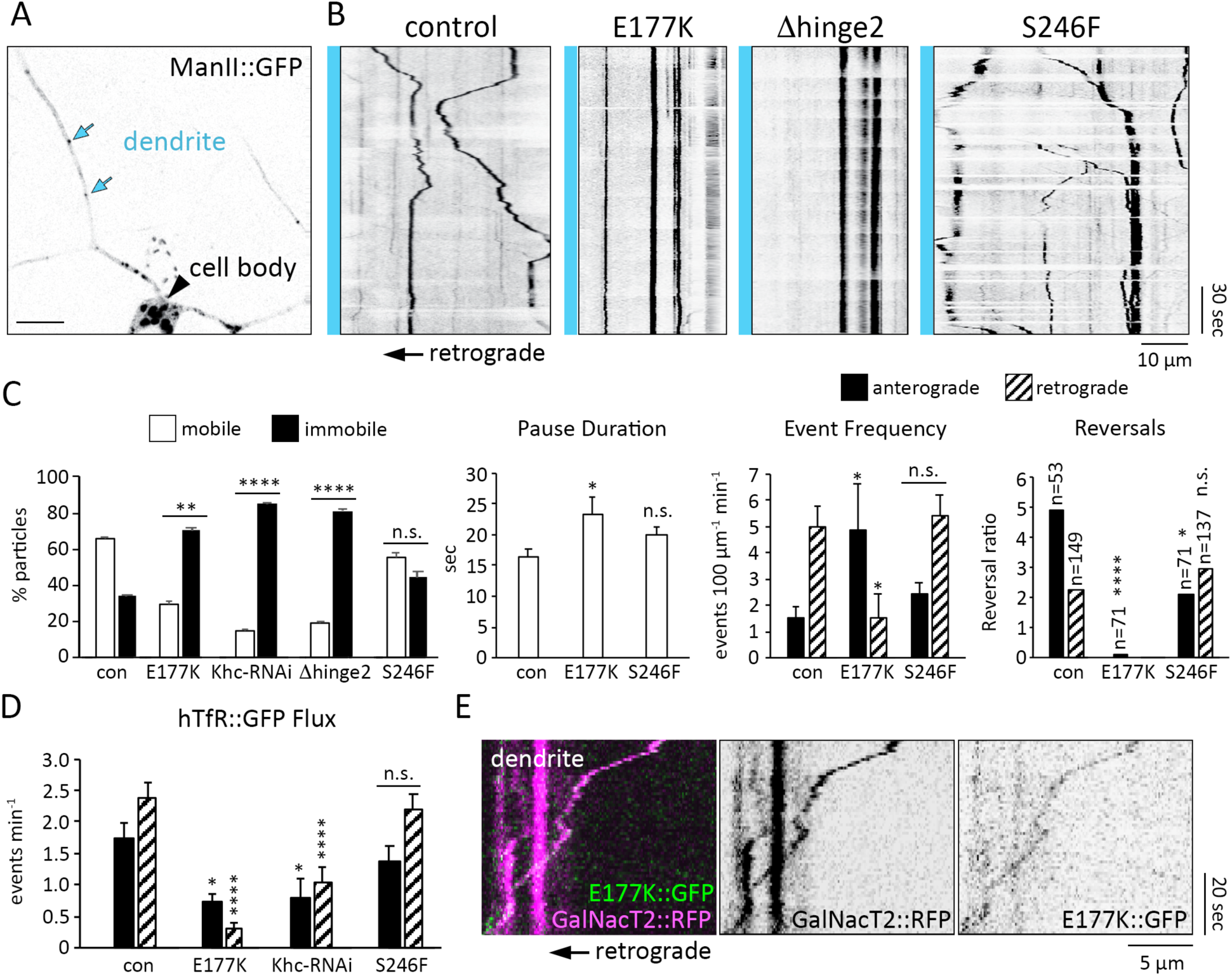
Khc autoinhibition mutants decrease the motility of Golgi outposts and hTfR-positive vesicles in dendrites. (A) Localization of Golgi outposts in control dendrites. The blue arrows highlight representative Golgi outposts used to create kymographs to analyze outpost dynamics. Scale bar = 10 μm. (B, C) Representative kymographs (B) and quantification (C) of Golgi outpost motility in control and mutant dendrites as indicated. Blue bar and arrow (bottom of kymograph) indicate the relative position of the cell body (left in all kymograph panels). Scale bars = 10 μm (x-axis) and 30 sec (y-axis). In control dendrites, robust movement of Golgi outposts in both directions is observed. In mutants with attenuated autoinhibition (*Khc* ^*E177K/-*^, *Khc* ^*Δhinge2/-*^), the majority of Golgi outposts are immobile (B and far left panel in C). The E177K mutation increases pause duration (middle left panel in C) and decreases the frequency of retrograde events (middle right panel in C). Anterograde-moving Golgi outposts are less likely to reverse direction after a pause in *Khc* ^*E177K/-*^ neurons versus control neurons (far right panel in C; the number of retrograde events was too small to include in the reversal ratio quantification). Diminishing Khc activity (*Khc* ^*S246F/-*^), does not significantly alter Golgi outpost motility (B and C). The percent mobile and immobile outposts in 9 (control, *Khc* ^*Δhinge2/-*^), 10 (*Khc* ^*E177K/-*^), and 11 (*Khc-RNAi* and *Khc* ^*S246F/-*^) neurons in at least five larvae were quantified. The outpost motility, event frequency, pause duration, and reversal ratio was calculated for 9 (control), 7 (*Khc* ^*E177K/-*^), and 10 (*Khc* ^*S246F/-*^) neurons in at least five larvae (the *Khc-RNAi* and *Khc* ^*Δhinge2/-*^ neurons had too few events for these calculations). The reversal ratio represents the number of events that reversed direction after a pause relative to the number of events that did not reverse direction; significant differences between the reversal ratios were determined via Chi-squared tests (n = number of events following a pause). (D) Quantification of hTfR::GFP flux in dendrites (average ± SEM) in 9 (*Khc-RNAi* and *Khc* ^*S246F/-*^), 11 (control), and 12 (*Khc* ^*E177K/-*^), neurons in at least four larvae. Flux is reduced in both directions in *Khc* ^*E177K/-*^ and *Khc-RNAi* neurons relative to controls, though retrograde flux is a more substantially reduced. *P=0.05–0.01, **P=0.01-0.001, ^****^P < 0.0001, and n.s. = not significant when compared to control neurons and evaluated by one-way ANOVA and Tukey post-hoc test (C, D). (E) Kymograph showing Khc-E177K::sfGFP (green) co-localizes with a Golgi outpost (GalNacT2::RFP, magenta) moving in a retrograde direction indicative of kinesin-mediated transport. Scale bars = 5 μm (x-axis) and 20 sec (y-axis).

Next, we attempted to visualize Golgi outposts being transported by the E177K mutant motor. We found it technically challenging to visualize Khc-E177K::sfGFP in mutant *Khc* ^*E177K::sfGFP11/-*^ neurons, but we did observe that Khc-E177K::sfGFP co-localized with retrogradely moving Golgi outposts in the dendrites of heterozygous *Khc* ^*E177K::sfGFP11/+*^ neurons (Fig. 7 E). Since one gene copy of the E177K mutation is not sufficient to drive outposts into axons, we could only observe co-localization in dendrites (only one Khc tail in a motor dimer is needed to inhibit activity). The co-localization between the E177K mutant motor and Golgi outposts in dendrites is consistent with our model that the E177K mutant motor is able to actively transport Golgi outposts into axons.

### Khc interacts with dynein to regulate Golgi outpost localization

Golgi outposts are transported into dendrites by dynein (Arthur et al., 2015; Ye et al., 2007; Zheng et al., 2008), and the dendrite-specific localization of Golgi outposts may depend on a regulated balance of dynein and kinesin-1 activities. We used genetic interaction tests to determine whether disrupting the motor balance to favor Khc results in the ectopic axonal localization of outposts. We used RNAi targeting the dynein light intermediate chain (dlic) subunit (*dlic-RNAi*) to reduce dynein function. A strong decrease in dynein function (*dlic-RNAi* and *dicer*) resulted in the ectopic accumulation of Golgi outposts at axon terminals, consistent with the idea that a shift favoring kinesin enables the motor to pull outposts into axons (Fig. 8 A). A weak dynein loss-of-function (*dlic-RNAi* alone) did not affect outpost distribution, providing a paradigm to test for genetic interactions (Fig. 8 A). In the weak dynein loss-of-function background, Golgi outpost distribution was not affected by reducing Khc activity by either removing one copy of *Khc* or introducing the Khc S246F mutation that decreases ATPase activity (Fig. 8, B, C and F). This is consistent with the notion that attenuating the activity of both dynein and kinesin-1 simultaneously maintains a motor balance that restricts Golgi outposts to dendrites.

**Figure 8:**
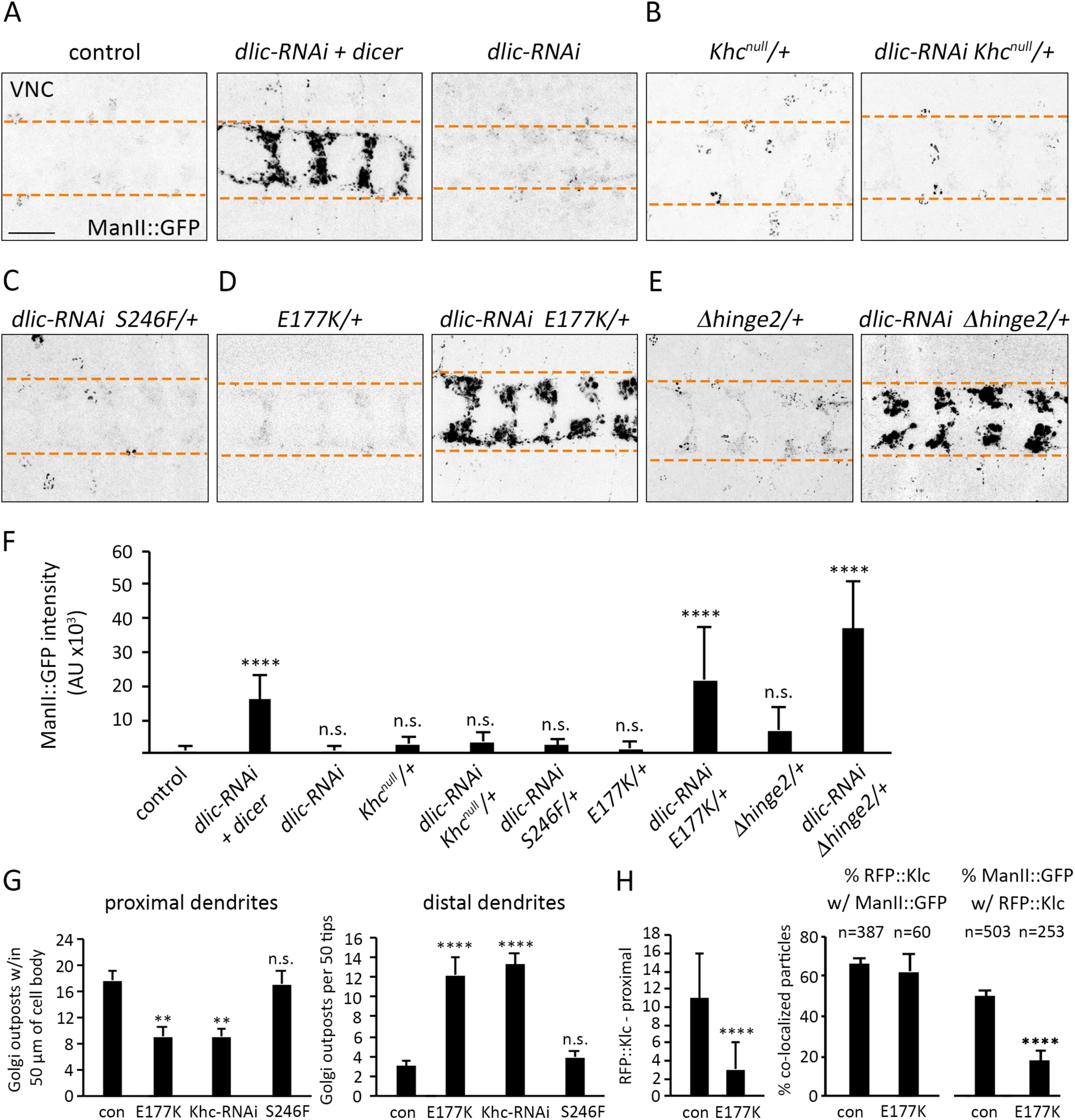
Khc autoinhibition mutants interact with dynein to regulate Golgi outpost localization. (A-F) Representative images (A-E) and quantification (F) of Golgi outposts at axon terminals in the VNCs of control and mutant animals as indicated. Scale bar = 25 μm. Dashed lines demarcate borders of axon terminals (A-E). One copy of *dlic-RNAi* by itself does not result in ectopic Golgi outposts in axon terminals but enhancing *dlic-RNAi* with dicer results in a substantial accumulation (A, F). Reducing *Khc* dosage (*Khc* ^*null/+*^) or activity (*Khc* ^*S*246/+^) does not enhance *dlic-RNAi*, and Golgi outposts remain excluded from axon terminals (B, C, F). In animals with one copy of the E177K or hinge2 mutation, Golgi outposts are absent from axon terminals; however, *Khc* ^*E177K/+*^ or *Khchinge2/+* combined with *dlic-RNAi* results in ectopic Golgi outposts in axon terminals (D-F). Quantification of ManII::GFP signal intensity (average ± SD) at axon terminals in 18 (*Khchinge2/+*), 19 (*Khc* ^*E177K/+*^ and *Khc* ^*null/+*^), 20 (*dlic-RNAi* + *dicer* and *dlic-RNAi* + *Khc* ^*S*246*F/+*^), 24 (*dlic-RNAi*), and 25 (*dlic-RNAi* + *Khc* ^*null/+*^, *dlic-RNAi* + *Khc* ^*E*177*K/+*^, and *dlic-RNAi* + *Khchinge2/+*) VNC segments in at least four animals (F). (G) Quantification of Golgi outpost distribution (average ± SEM) in proximal (left) and distal (right) regions of class IV sensory dendrite arbors. Proximal: Outposts within 50 μm of the cell body were counted in 14 (*Khc* ^*E177K/-*^), 15 (control, *Khc-RNAi*, and *Khc* ^*S246F/-*^), and 16 (*Khc* ^*E177K,S246F/-*^) neurons in at least six larvae. Distal: Outposts in a 125 × 250 μm area encompassing the distal arbors of 20 (*Khc-RNAi, Khc* ^*S246F/-*^, and *Khc* ^*E177K,S^246^F/-*^) and 24 (control and *Khc* ^*E177K/-*^) neurons in at least six larvae were counted and normalized to the number of dendrite tips (control dendrite arbors have ∼50 dendrite tips in the distal region). In both the *Khc* ^*E177K/-*^ and *Khc-RNAi* dendrite arbors, the number of proximal Golgi outposts decreases at the same time that the number of distal outposts increases. (H) Quantification of RFP::Klc puncta within 50 μm of the cell body (proximal) in control and E177K mutant neurons (left) and co-localization of RFP::Klc and ManII::GFP in proximal dendrites of control and *Khc* ^*E177K/-*^ neurons (right). The number of RFP::Klc puncta is reduced, but the percentage of RFP::Klc puncta that co-localize with ManII::GFP is unchanged in *Khc* ^*E177K/-*^ neurons, which suggests the association between Klc and outposts is not affected by the E177K mutation. The number of RFP::Klc-positive Golgi outposts is reduced, reflective of the overall decrease in RFP::Klc in dendrites. ^*^P=0.05–0.01, ^**^P=0.01-0.001, ^****^P < 0.0001, and n.s. = not significant in comparison to control and evaluated by one-way ANOVA with Tukey post-hoc test (F-H).

Next, we tested whether reducing Khc autoinhibition in the weak dynein loss-of-function background would disrupt the motor balance and Golgi outpost localization. One copy of *Khc* ^*E177K*^ in a control background had no effect on Golgi outpost distribution (Fig. 8, D and F). However, one copy of *Khc* ^*E*^177^*K*^ coupled with a mild reduction of dynein function resulted in the mislocalization of Golgi outposts to axon terminals (Fig. 8, D and F). Similarly, one copy of *Khchinge2* by itself was insufficient to cause the mislocalization of Golgi outposts, but one copy of *Khchinge2* combined with *dlic-RNAi* resulted in ectopic Golgi outposts at axon terminals (Fig. 8, E and F). These results support the idea that disrupting Khc autoinhibition tips the dynein-kinesin balance towards kinesin-1 and transport into axons.

If a balance of dynein and kinesin-1 regulates Golgi outpost distribution, then reducing kinesin-1 levels in dendrites should shift the distribution of outposts distally in the arbor. Consistent with this idea, knocking-down Khc reduced the number of Golgi outposts proximally and increased outposts distally (Fig. 8 G). The E177K mutation, which significantly reduces Khc levels in dendrites, resulted in a similar proximal decrease and distal increase in outposts. Although there are more Golgi outposts distally, we did not observe a corresponding shift in dendrite branches such as has been previously reported for mutations that alter outpost distribution or the dynein-kinesin balance (Arthur et al., 2015; Satoh et al., 2008; Taylor et al., 2015; Ye et al., 2007; Zheng et al., 2008). It is possible that kinesin-1 acts downstream of Golgi outposts to promote dendrite branching, or that Khc disrupts the transport of additional factors needed for branching. Of the Golgi outposts that remained in the proximal dendritic arbor of the E177K mutants, significantly fewer co-localized with RFP::Klc and overall fewer RFP::Klc puncta were present, paralleling the reduction in Khc levels and Golgi outpost motility (Fig. 8 H).

## Discussion

Neuronal function relies on the accurate delivery of proteins, vesicles, and organelles to axons and dendrites by molecular motors. Our in vivo structure-function analysis of endogenous Khc, combined with in vitro single-molecule experiments, indicate that regulated autoinhibition of kinesin-1 is necessary for the polarized dendritic distribution of Golgi outposts and hTfR-positive vesicles. Previous studies suggested that autoinhibition may be dispensable for some Khc-mediated activities while the autoinhibitory IAK tail motif is critical to axonal transport (Moua et al., 2011; Williams et al., 2014). Our results indicate that kinesin-1 autoinhibition has a previously unrecognized function in preventing the motor from carrying Golgi outposts into axons. Relieving autoinhibition enhances Khc activity and drives the cargo-bound motor into axons, depleting dendrites of Khc and perturbing dendrite arborization. Disrupting autoinhibition generates phenotypes that resemble kinesin-1 loss-of-function, but these effects can be reversed by decreasing ATPase activity, indicating they are due to a gain in motor activity, not a loss. It is likely that the depletion of the motor from a particular cellular compartment, such as dendrites, contributes to the loss-of-function phenotypes that we and others have observed when kinesin-1 autoinhibition is disrupted. Thus, tight regulation of kinesin-1 activity is critical to the polarized localization of dendritic cargo, the motor itself, and neuronal morphogenesis.

We had initially sought to determine whether kinesin-1 is stopped from carrying dendritic cargo into axons by reading out microtubule-based signals. In the kinesin-1 microtubule-binding domain, three loops have been implicated in recognizing microtubule-based signals, such as microtubule post-translational modifications and/or GTP-tubulin subunits, that differ between axons and dendrites (Bentley and Banker, 2016). Our studies reveal that loop 11 of the kinesin-1 microtubule-binding domain can be replaced by that of the kinesin-3 motor without affecting animal survival. However, the endogenous loop 12 and 5-loop 8 sequences are required for normal kinesin-1 function. Swapping loop 12 may decrease animal survival due to the K-loop in loop 12 of kinesin-3 increasing the on-rate of the motor onto microtubules (Soppina and Verhey, 2014). Our work indicates that the effects of mutating residues in 5-loop 8 are not due to altered microtubule track selection, but rather because the mutant kinesin-1 is no longer regulated by autoinhibition.

Within loop 8, we found that an essential short sequence, TERF, which differs between kinesin-1 and kinesin-3, is highly sensitive to the nature of the substitution. While changing the glutamate residue to glutamine or alanine had no effect on animal survival, the charge-reversal substitution of glutamate with lysine or arginine resulted in animal lethality and altered dendritic trafficking. Based on previous studies, we expected that the underlying mechanism would be either an alteration in kinesin-1 responsiveness to microtubule-based signals, such as microtubule detyrosination (Huang and Banker, 2012; Konishi and Setou, 2009), or a disruption of autoinhibition (Kaan et al., 2011). We found that the E177K mutation increases motor activity in vitro, causes the full-length motor to localize to axon terminals, and can be combined with dynein loss-of-function to drive outposts into axons. Additionally, autoinhibition-disrupting mutations in the Khc tail domain also result in the axonal mislocalization of Golgi outposts. We thus favor the idea that it is the loss of Khc inhibition that is responsible for altered dendritic trafficking in E177K mutant flies. While we cannot rule out the possibility that E177 is involved in track selection, it is notable that structural studies indicate the kinesin-1 TERF sequence does not directly interact with tubulin and thus may not directly respond to microtubule modifications (Atherton et al., 2014; Gigant et al., 2013). Combined, our data suggest that a key function of E177 is to regulate motor activity through autoinhibition, which we propose functions as a brake on the axonal localization of the motor when it is carrying dendritic cargo. It will be interesting to test the effects of E177K mutation and loss of autoinhibition on the localization and function of full-length kinesin-1 in mammalian neurons, whose dendritic microtubules have a more mixed polarity than fly.

Our autoinhibition model does not exclude other models of polarized transport, and indeed we believe that the regulated autoinhibition of kinesin-1 acts in concert with other mechanisms to regulate cargo distribution. Furthermore, models that take into account potential indirect effects of kinesin-1 autoinhibition on outpost localization are also possible. Kinesin-1 has been previously implicated in localizing dynein in axons and dendrites and some aspects of dynein activity may depend on kinesin-1 (Hancock, 2014; Satoh et al., 2008; Twelvetrees et al., 2016). Interfering with kinesin-1 autoinhibition might disrupt the localization and/or function of dynein and thereby enable a kinesin to pull outposts into axons. While this kinesin could be kinesin-1 itself, it is also possible that an additional kinesin is involved in transporting outposts. The best candidates for such another kinesin would be kinesin-2 or kinesin-3, which co-transport some cargo with kinesin-1 (Guardia et al., 2016; Gumy et al., 2017; Hendricks et al., 2010; Kulkarni et al., 2017; Lim et al., 2017; Ruane et al., 2016).

Our studies raise the question of how kinesin-1 autoinhibition is spatially regulated. It is likely that the autoinhibition of kinesin-1 attached to dendritic cargo would be triggered in the cell body or proximal axon, which is thought to function as a barrier to the entry of dendritic cargo and Golgi in invertebrate and vertebrate neurons (Bentley and Banker, 2016; Edwards et al., 2013; Rolls and Jegla, 2015). In the proximal axon of fly sensory neurons, Golgi outposts co-localize with the dynein co-factor NudE, which likely functions with dynein to localize outposts to dendrites (Arthur et al., 2015; Lin et al., 2015; Ye et al., 2007; Zheng et al., 2008). The mammalian NudE ortholog Ndel1 is proposed to “catch” dendritic cargos and prime them for dynein-mediated transport into dendrites (Klinman et al., 2017; Kuijpers et al., 2016). We found that simultaneously decreasing dynein activity and kinesin-1 autoinhibition resulted in ectopic axonal Golgi outposts. This suggests that the selective transport of Golgi outposts to dendrites relies on coordinating the activation of a dendrite-targeted motor (dynein) and the inhibition of a motor that targets axons (kinesin). A key future goal is to determine how kinesin-1 autoinhibition is spatially regulated to mediate the proper distribution of cargo to axons or dendrites to maintain neuronal polarity.

## Materials and methods

### Molecular cloning

*Khc* knock-in alleles: The *Khc* knock-in alleles were generated using the fly strain *Khc* ^*KO-attP*^, in which *Khc* is knocked-out and replaced with an *attP* site for rapid site-directed integration of new alleles (Winding et al., 2016). To knock-in new Khc alleles, an integration plasmid (*pGE-attB-GMR*) (Huang et al., 2009) containing the new allele was injected into *Khc* ^*KO-attP*^ embryos and integrated via ϕC31-mediated recombination. The following mutations were introduced by PCR-based mutagenesis using Q5^®^ High-fidelity polymerase (NEB): loop-11 swap, loop-12 swap, SKLA, SQLA, E177A, E177K, E177R, hinge2, R947E, K944E, S246F, E177K + S246F, S181D +S182D, and S181A + S182A. To make the β5-loop-8 swap mutant, the *unc-104* sequence that encodes 5-loop 8 was ordered as a gBlock (IDT) and the final construct was created via Gibson Assembly (NEB). The amino acid sequences that were swapped are as follows: Khc 5-loop 8 (KIRDLLDVSKVNLSVHEDKNRVPYVKGATERFVSS) was substituted with RVRDLLNPKNKGNLRVREHPLLGPYVEDLSKLAVTD from unc-104; Khc loop 11 (KVSKTGAEGTVLDEAK) was substituted with RADSTGAKGTRLKEGA from unc-104; Khc loop 12 (GNKT) was substituted with VASKKKNTKKAD from unc-104. To make Khc::sfGFP11, the sfGFP11 peptide (multimerized 7X) was added to the C-terminal end of Khc with a GGSGG linker between Khc and sfGFP11 and successive sfGFP11 peptides. All knock-in plasmids were fully sequenced before injection (BestGene Inc.).

Transgenic flies: *UAS-Khc(1-401)::sfGFP* was assembled by first using HiFi DNA assembly (NEB) to piece together a superfolder GFP (sfGFP)-encoding gene block (IDT) and *pIHEU-MCS* plasmid (plasmid #58375, Addgene) to create *sfGFP-pIHEU*. The *pTTSTOP* plasmid containing truncated Khc, residues 1-401, (plasmid #41751, Addgene; (Thoresen and Gelles, 2008)) was digested, subcloned, and then ligated into *sfGFP-pIHEU* to create *UAS-Khc(1-401)::sfGFP*. PCR-based mutagenesis of *UAS-Khc(1-401)::sfGFP* using Q5® High-fidelity polymerase (NEB) and Gibson Assembly created *UAS-Khc(1-401)-E177K::sfGFP*. The plasmids were injected into strain *P{CaryP}attP40* (BestGene Inc.). *Ppk-ManII::GFP*, which expresses ManII::GFP (Ye et al., 2007) under the control of the *ppk* enhancer, was created by PCR-amplifying *ManII::GFP* and cloning it into *pDEST-APPHIH*, which contains a 1-kb *ppk* enhancer fragment (Chun Han, Cornell University). *Ppk-ManII::GFP* was injected into strain *PBac{yellow[+]-attP-3B}VK00037* (BDSC stock #9752, BestGene Inc.). *UAS-sfGFP(1-10)*, which was made by cloning sfGFP(1-10) into *pACUH* (plasmid #58374, Addgene) (Kamiyama et al., 2016), was injected into strain *PBac{y[+]-attP-3B}VK00027* (Rainbow Transgenic Flies, Inc.).

### Fly genetics

The Khc alleles *Khc* ^*27*^(null allele), *Khc* ^*22*^ (amino acid replacement P945S), and *Khc* ^*23*^ (amino acid replacement E164K) were provided by W. Saxton, University of California, Santa Cruz, Santa Cruz, CA (Brendza et al., 1999). UAS-Khc::BFP was provided by V. Gelfand, Northwestern University, Chicago, IL (Winding et al., 2016). The following fly strains were obtained from the Bloomington Drosophila Stock Center (BDSC, Bloomington, IN) and the Vienna Drosophila Resource Center (VDRC, Vienna, Austria): *UAS-hTfR::GFP* (BDSC stocks #36858), *ppk-CD4::Tomato* (BDSC stock #35845), *ppk-CD4::GFP* (BDSC stock #35843), *ppk-Gal4* (BDSC stock #32078 and 32079), *ensconsin* alleles, including *ensc* ^*Δ*^ *C*, which lacks the majority of coding exons (Sung et al., 2008) (BDSC stock #51318), and *Df(3L)ensc* ^*Δ*^ *3277*, a deficiency that removes *ensconsin* (BDSC stock #51319), *UAS-Khc-RNAi* (BDSC stock #35770), *UAS-dicer2* (BDSC stock #24650), and *UAS-dlic-RNAi* (VDRC stock #41686). All transgenic markers (e.g. *ppk-ManII::GFP*) were used as a single copy. Khc mutants were analyzed in trans to the null allele *Khc* ^*27*^ unless otherwise noted.

### Image capture and analysis

All neuron imaging was carried out on a Leica SP5 confocal microscope. The dorsal class IV dendritic arborization neurons (ddaC) in the abdominal segments of control and mutant larvae were imaged in live animals. Live larvae were mounted on slides in a solution of 50% PBS and 50% glycerol and immobilized by pressing a cover slip mounted on top of two lines of vacuum grease spacers flanking the animal.

To quantify the number of ManII-positive Golgi outposts in axons, third-instar larvae expressing ManII::GFP and CD4::Tomato in ddaC neurons were imaged at a resolution of 1024×1024 pixels using a 40×1.3 NA oil-immersion objective. Z-stacks 10-20 μm thick (1.5 μm per z-step) were max projected and the number of ManII::GFP puncta was quantified using the FIJI particle analysis function. The ManII::GFP signal in black-and-white images was inverted and subject to threshold with a cut-off grey value of 170. ManII::GFP-positive puncta between 0.15-15.00 μm in diameter within 75 μm of the cell body (axon) or a 50 μm radius of the cell body (proximal dendrites) were counted. To calculate Golgi outposts per dendrite tip, a box 125 μm × 250 μm was drawn around the distal tips closest to the dorsal midline. This area was inverted and subject to threshold with a cut-off grey value of 140. ManII::GFP-positive puncta between 0.10-15.00 μm in diameter were counted. The number of tips in the same region was counted using the CD4::Tomato channel.

Golgi outpost co-localization with Klc was determined by simultaneous imaging of ManII::GFP and RFP::Klc for 2-5 minutes in ddaC dendrites of third instar larvae. Movies were captured at a rate of 0.667 frames/second and a resolution of 1024×512 pixels using a 40×1.3 NA oil-immersion objective. Movies were stabilized using the FIJI Image Stabilizer plugin and kymographs were generated using ImageJ. Kymographs were used to determine co-localization between Golgi outposts and Klc as well as to calculate the directionality and velocity.

Golgi outpost motility was quantified by imaging ManII::GFP puncta for 4–6 minutes in one or two ddaC neurons per larvae at 72 hours (h) after egg laying (AEL). Movies were captured at a rate of 0.895 frames/second and a resolution of 1024×512 pixels using a 40×1.3 NA oil-immersion objective. ManII::GFP puncta within 100 μm of the cell body in primary and secondary dendrites were analyzed. Movies were stabilized using the FIJI Image Stabilizer plugin and kymographs were generated using Metamorph. Position and time data from Metamorph were exported to Excel to quantify directionality. Puncta that traveled consistently in one direction were classified as anterograde or retrograde, whereas those that switched direction were classified as bidirectional. Puncta that did not move during the movie were classified as paused. Kymographs were further in analyzed Metamorph to quantify the number, frequency, velocity, run length, and pause duration of anterograde and retrograde events. The event frequency was calculated only for movies with motile outposts. Particles that moved, paused for at least 3 frames (2.62 sec) and then resumed movement were used to calculate direction changes and determine the frequency of direction switching.

Dendrite length, branch points, and Sholl analysis were quantified by imaging 2-4 ddaC neurons from segments A3-A6 in larvae at 72 and 96 h AEL at a resolution of 2048×2048 pixels using a 20×0.7 NA oil immersion objective. Z-stacks 10-15-μm thick (1 μm per z-step) were analyzed using the FilamentTracer module in the Imaris software program to obtain dendrite length, branch point number, and Sholl crossings. Neurons were analyzed individually by masking signals from neighboring neurons. Data was exported to Excel for statistical analysis. Axon branching was assessed for late third instar larvae by counting branch points in individual axons. Axons were split into three categories: unbranched, less than five branches, or five and greater branches.

Truncated Khc(1-401)::sfGFP localization was quantified by imaging 2-4 ddaC neurons from segments A3-A5 in third instar larvae at a resolution of 1024 × 1024 pixels using a 40×1.3 NA oil immersion objective. Z-stacks 10-15-μm thick (1 μm per z-step) were analyzed using FIJI software to determine the background-subtracted GFP intensity and Tomato signal in the proximal dendrites and axons 50 μm from the cell body. The ratio of GFP/Tomato was calculated for each neurite analyzed.

Microtubule polarity was analyzed by imaging EB1::GFP puncta within 100 μm of the cell body for 4–6 minutes in one or two ddaC neurons per larvae at 72 h AEL. Movies were captured at a rate of 0.895 frames/second and a resolution of 1024×512 pixels using a 40×1.3 NA oil-immersion objective. Movies were stabilized using the FIJI Image Stabilizer plugin and kymographs were generated using Metamorph. Position and time data from Metamorph were exported to Excel to quantify directionality.

To image ManII::GFP, Khc(1-401)::sfGFP, and hTfR::GFP in the VNC, brains from 3^rd^ instar larvae were dissected in 1X PBS, fixed in 4% paraformaldehyde for 20 minutes, washed with 1X PBS, and mounted on slides in a solution of elvanol between vacuum grease spacers. Fixed VNCs were imaged using a 40×1.3 NA oil-immersion objective. Z-stacks 20-30 μm thick (1 μm per z-step) were analyzed using FIJI. The background-subtracted, integrated fluorescence intensity within a 20 × 50 μm box drawn around commissures was quantified in 4-5 terminals per larvae (segments A2-A6).

The fluorescent signal of reconstituted sfGFP in axon terminals expressing Khc::sfGFP11 and sfGFP(1-10) was quantified by determining the background-subtracted, integrated fluorescence intensity of both Khc::sfGFP and CD4::Tomato within a 20 × 50 μm box drawn around commissures (segments A2-A6). The ratio of GFP/Tomato signal was determined for each commissure analyzed. The Khc::sfGFP/Tomato signal in dendrites was similarly quantified in FIJI by drawing 50 μm-long line starting from the cell body and tracing the primary dendrite with the brightest GFP signal. The average intensity over 10-μm segments was statistically analyzed.

### Antibody Staining

To image bruchpilot (Brp), cysteine string protein, and synapsin (syn) signal in sensory neurons, 3^rd^ instar larvae were dissected in 1X PBS, fixed in 4% paraformaldehyde for 45 minutes, washed with 1X PBS, permeabilized in 1X PBS, 0.3% Triton X-100 (PBSTx) for 20 min, washed with PBSTx, quenched in 50 mM NH_4_Cl for 10 min, washed with PBSTx, and incubated in block for at least an hour. Larval fillets were incubated in primary antibody overnight, washed (3 × 30 min in PBSTx), and then incubated in fluorescent secondary antibodies overnight. Fillets were then washed in PBSTx (3 × 30 min) and mounted in a solution of elvanol. Fixed larvae were imaged using a 40×1.3 NA oil-immersion objective. Z-stacks 20-30 μm thick (1 μm per z-step) were analyzed using FIJI. Antibodies to Brp (nc82), cysteine string protein (DCSP-2 6D6), and synapsin (3C11) were obtained from the Developmental Studies Hybridoma Bank.

### S2 cell lysates

Drosophila S2 cells were cultured in Schneider’s Drosophila medium (Gibco) supplemented with 10% (vol/vol) fetal bovine serum (FBS, Hyclone) at 25 °C. Drosophila Khc constructs in the vector pMT were transfected into cells using Cellfectin II (Invitrogen) and protein expression was induced by adding 1 mM CuSO_4_ to the medium 4-5 h post-transfection. The cells were harvested after 48 h.

Motor-expressing S2 cells were harvested and centrifuged at low-speed at 4 °C. The cell pellet was washed once with PBS and resuspended in ice-cold lysis buffer (25 mM HEPES/KOH, pH 7.4, 115 mM potassium acetate, 5 mM sodium acetate, 5 mM MgCl_2_, 0.5 mM EGTA, 1% vol/vol Triton X-100) freshly supplemented with 1 mM ATP, 1 mM phenylmethylsulphonyl fluoride (PMSF), protease inhibitors cocktail (P8340, Sigma), 10% glycerol and 1 mM DTT. After clarifying the lysate by centrifugation at 16,000g at 4 °C, aliquots were snap frozen in liquid nitrogen and stored at −80°C until further use. The relative amount of motor across lysates was calculated by a dot-blot in which serial dilutions of each S2 lysate were spotted onto a nitrocellulose membrane (GE Healthcare Life Science). The membrane was air dried and immunoblotted with a polyclonal antibody to kinesin-1 generated against the motor domain peptide (KLSGKLYLVDLAGSEKVSKTGAEG) (Verhey et al., 1998). The amount of Khc in each spot was quantified (Image J) and spots within the linear regime were used to normalize motor levels for addition to single-molecule motility assays.

### Single-molecule motility assays

All single-molecule assays were performed at room temperature in a flow cell (∼10 μl volume) prepared by attaching a clean #1.5 coverslip to a glass slide with double-sided tape. HiLyte-647-labeled microtubules were polymerized from a mixture of HiLy647-labeled and unlabeled tubulins (Cytoskeleton) in BRB80 buffer (80 mM PIPES/KOH, pH 6.8, 1 mM MgCl_2_, 1mM EGTA) supplemented with 1 mM GTP at 37°C for 15 min. After addition of five volumes of prewarmed BRB80 containing 20 μM taxol and an additional 15 min incubation at 37°C, polymerized microtubules were stored at room temperature in the dark for further use. Polymerized microtubules were diluted in BRB80 Buffer containing 10 μM taxol and then infused into flow-cells and incubated for 5 min at room temperature for non-specific adsorption to the coverslip. Subsequently, Blocking Buffer [1 mg/ml casein in P12 buffer (12 mM PIPES/KOH, pH 6.8, 2 mM MgCl_2_, 1mM EGTA) with 10 μM taxol] was infused and incubated for another 5 min. Finally, kinesin motors in the motility mixture [0.5-1μl of S2 cell lysate, 2 mM ATP, 3 mg/ml casein, 10 μM taxol, and oxygen-scavenging (1mM DTT, 1mM MgCl_2_, 10 mM glucose, and 0.2 mg/ml glucose oxidase, 0.08 mg/ml catalase) in P12 buffer] was added to the flow cell. The flow cells were sealed with molten paraffin wax and imaged by TIRF microscopy using an inverted microscope Ti-E/B (Nikon) equipped with the perfect focus system (PFS) (Nikon), a 100X 1.49 N.A. oil immersion TIRF objective (Nikon), three 20 mW diode lasers (488 nm, 561 nm and 640 nm) and EMCCD detector (iXon X3DU897, Andor). Imaging of full length motors was carried out by continuous imaging at 200 ms per frame for 1 min and imaging of truncated motors was carried out by continuous imaging at 100 ms per frame for 30 s. Image acquisition was controlled by Nikon Elements software.

### Analysis of single-molecule motility data

Maximum intensity projections were generated from the movies and kymographs were produced by drawing along motor-decorated tracks (width= 3 pixels) using Nikon Elements software. Run length was calculated by taking the distance moved along the x-axis of the kymograph. Dwell time was calculated by taking the time of the event along the y-axis of the kymograph. Velocity was defined as the distance on the x axis of the kymograph divided by the time on the y axis of the kymograph. Full-length motors frequently paused during motility events and thus only the run length and velocity between pauses were analyzed. Motility events which started before image acquisition or finished after image acquisition were included in the analysis and thus the motility parameters for motors with long dwell times are an underestimate of the true values. Motility data from at least two independent experiments were pooled and analyzed.

Relative landing rates were calculated for truncated motors by normalizing motor protein levels across lysates to ensure equal motor input. The relative landing rate was defined as the number of events per unit microtubule length per unit time and are reported as # events (μm s)^−1^ ± SEM. Only events that lasted at least 5 frames (500 ms) were counted. Over 10 different microtubules from at least two independent experiments were analyzed for each construct.

### Statistical analysis

Data were analyzed in Stata software using Student’s unpaired *t-test*s to compare individual samples and one-way ANOVA and Tukey post-hoc test (a=0.01 was used to determine whether a significant difference existed) and *t-test*s post hoc to compare multiple samples. Significance levels are represented as follows: n.s. (not significant), *P=0.05–0.01, ^**^P=0.01–0.001, ^***^P=0.001–0.0001, and ^****^P<0.0001.

## Acknowledgments

We thank William Saxton (University of California, Santa Cruz, CA), Vladimir Gelfand (Northwestern University), the Bloomington Drosophila Stock Center, and the Vienna Drosophila Stock Center for supplying fly strains; Kevin Eliceiri and the Laboratory for Optical and Computational Instrumentation (LOCI, University of Wisconsin, Madison, WI) for image analysis support; and Lindsay Mosher and Dena Johnson-Schlitz for technical assistance. We thank Ivan Rayment (University of Wisconsin-Madison) and members of the Wildonger lab for insightful, enthusiastic discussions and comments on the manuscript.

This work was supported by start-up funds from the University of Wisconsin-Madison (J.W.) and grants from the National Institutes of Health (R21MH101688 and R21EB022798 to B.H., R01GM070862 to K.J.V., and R01NS102385 to J.W.).

---

**Figure S1:**
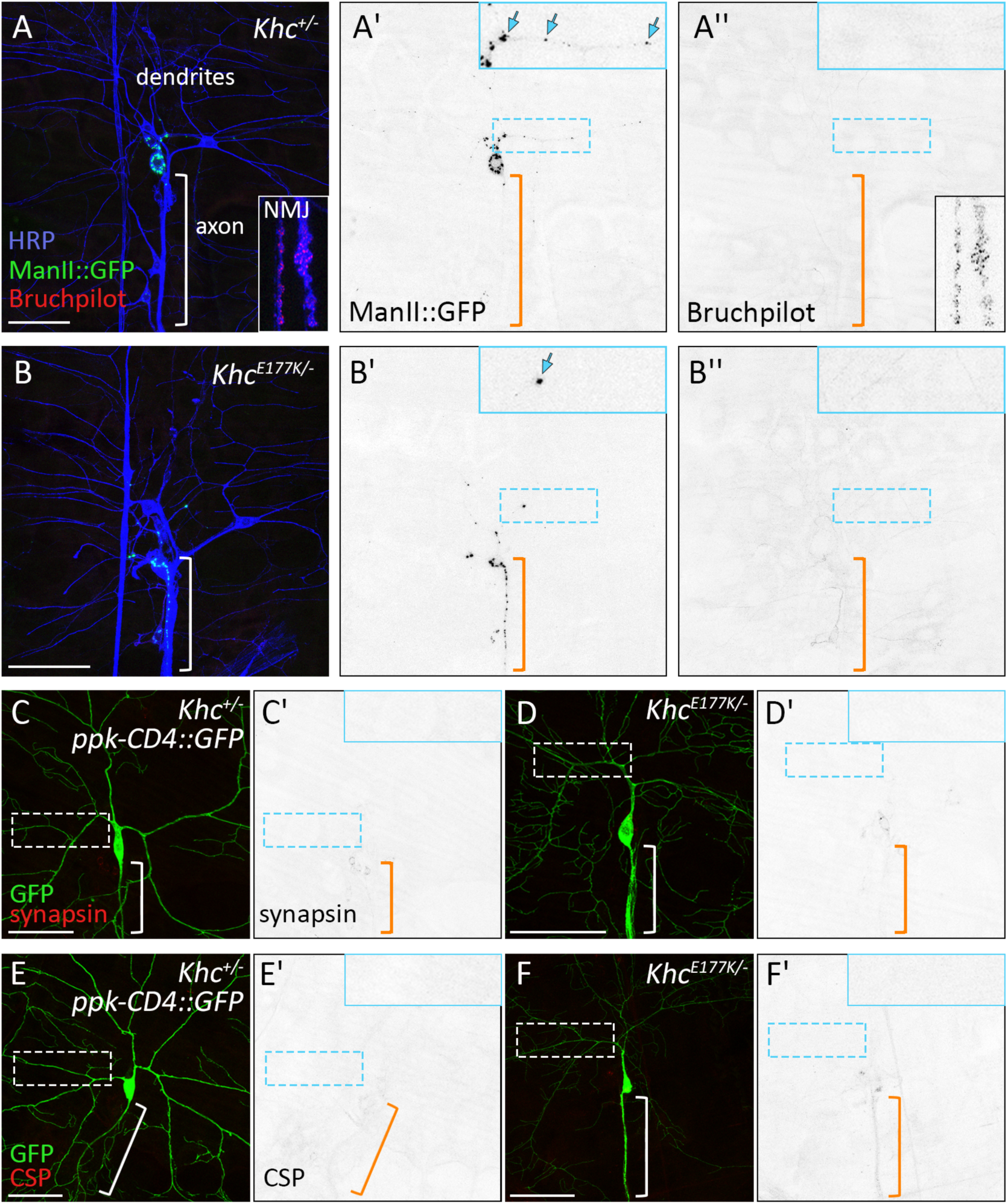
Localization of Brp, synapsin, and CSP in *Khc^+/-^* and *Khd^177K/-^* neurons. (A-F) Brp, synapsin, and CSP are absent from dendrites of *Khc^*A^* (A-A″,C, C^1^, E, and E^1^) and *Khd^U7K/^*-neurons (B-B″, D, D′, F, and F′). Insets (A′, A″, B′, B″_;_ C′, D′, E′, and F′) show zoomed view of dendrites, which are devoid of synaptic proteins but contain Golgi outposts (arrows in A′ and B^1^). *Khd^l77K/^*′ axons contain mislocalized Manll::GFP-positive Golgi outposts (B and B^1^). Inset in A and A″ shows Brp at a neuromuscular junction (NMJ), an internal staining control. Neuronal membranes are labeled by anti-horseradish peroxidase (A-B″) and CD4::GFP (C-F′).

**Figure S2:**
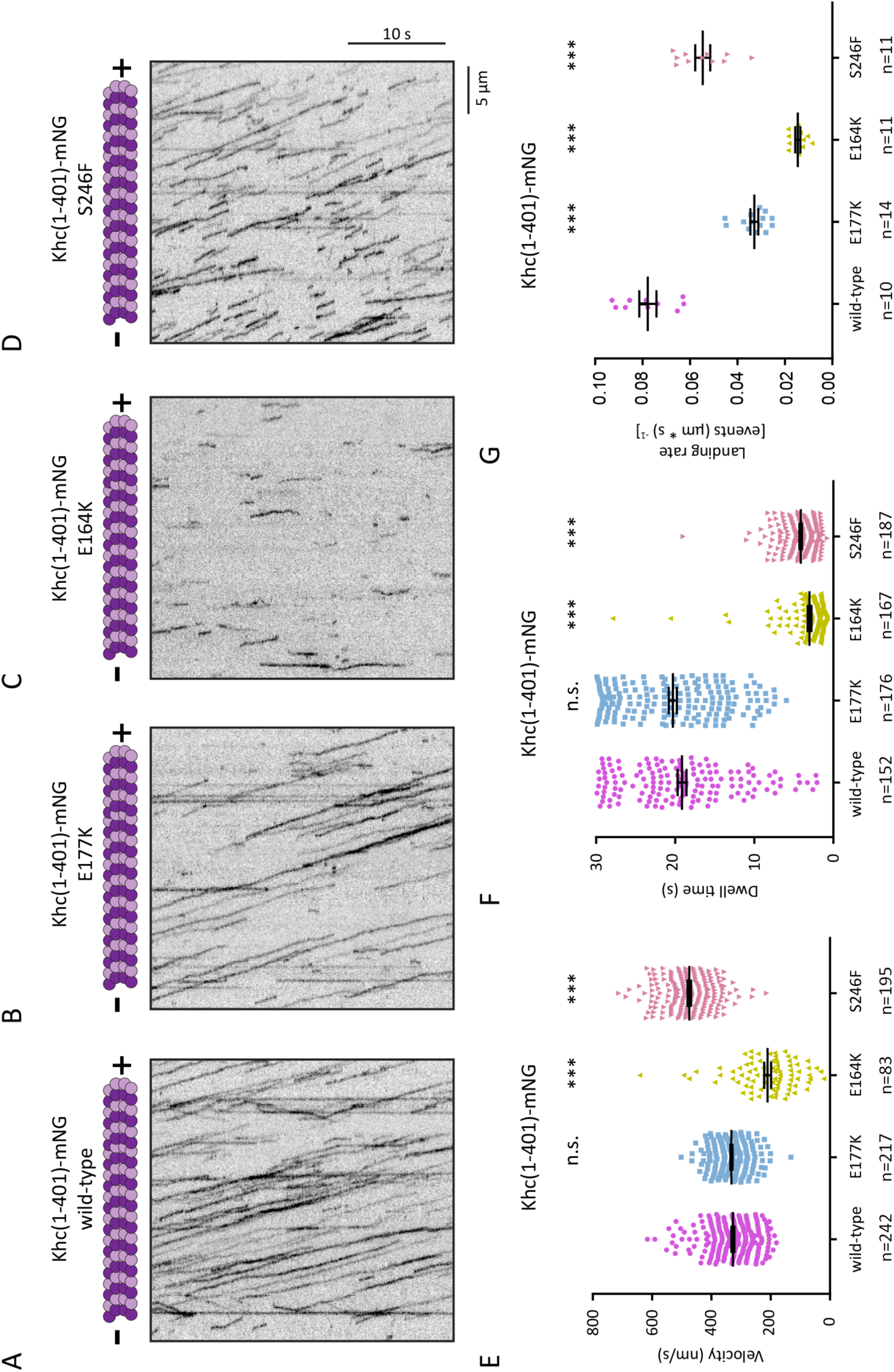
Effects of mutations on motility of truncated Khc. (A-D) Representative kymographs showing single-molecule motility of truncated wild-type KHC(1-401) and the indicated mutant versions. All motors were tagged with mNG at their C-terminus. Distance is on the x-axis and time is on the y-axis. Scale bars = 5μ.m (x-axis) and 10 s (y-axis). (E-G) Quantification of (E) velocities, (F) dwell time and (G) landing rate for each population of motors. Note that the truncated Khc(1-401)-S246F::mNG moves faster than wild-type, but the full-length S246F mutant motor moves more slowly (see Fig. 4). Scatterplots display Mean ± SEM from at least two independent experiments. ***p<0.01 and n,s. as compared to the wild-type motor (calculated by two-tailed t test).

**Figure S3:**
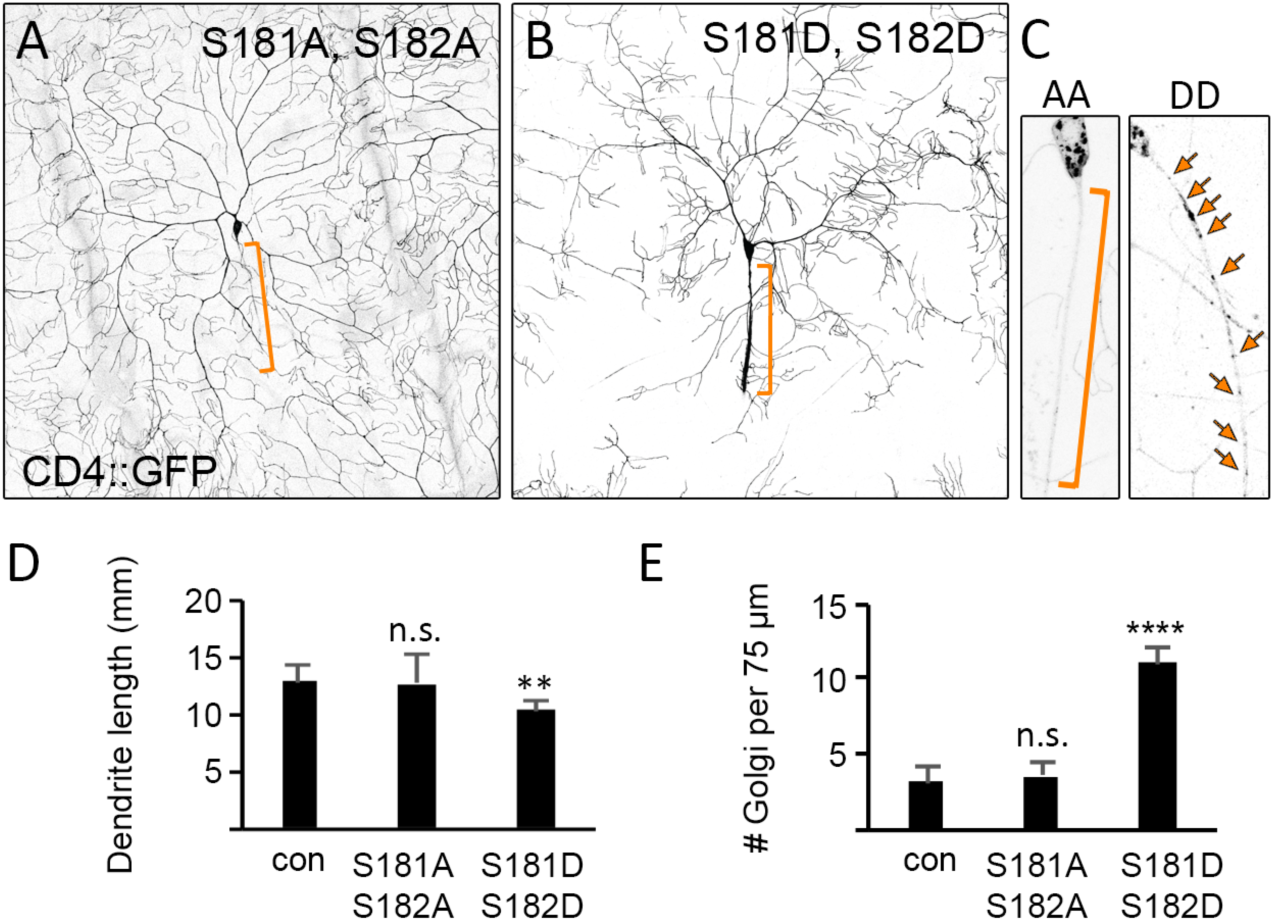
Effects of Khc phosphomutants on dendrite morphogenesis and Golgi outpost localization. (A,B,D) Dendrite arborization is not affected in *Khc*^5181A,5132AI^ neurons, but is reduced in the phosphomimetic *Khc*^51310,513201^ mutants. {C,E} Golgi outposts labeled by Manll::GFP are dendrite-specific in *Khc*^5181A,sis^2AI neurons (left), but localize ectopically to axons in *Khc*^51810,518201^ neurons (right). (*Khc*^51s1A,s1s2A/-^), and 8 (*Khc*^51s10,s1 s201^) neurons (E). ^**^P < 0.01, Quantification of dendrite length (average± SD} of 16 (control), 9 (average± SD} in the proximal 75 μm of axons in 9 (control), 8 ^****^P < 0.0001, n.s. = not significant in comparison to control and evaluated by one-way ANOVA and Tukey post-hoc test.

**Figure S4:**
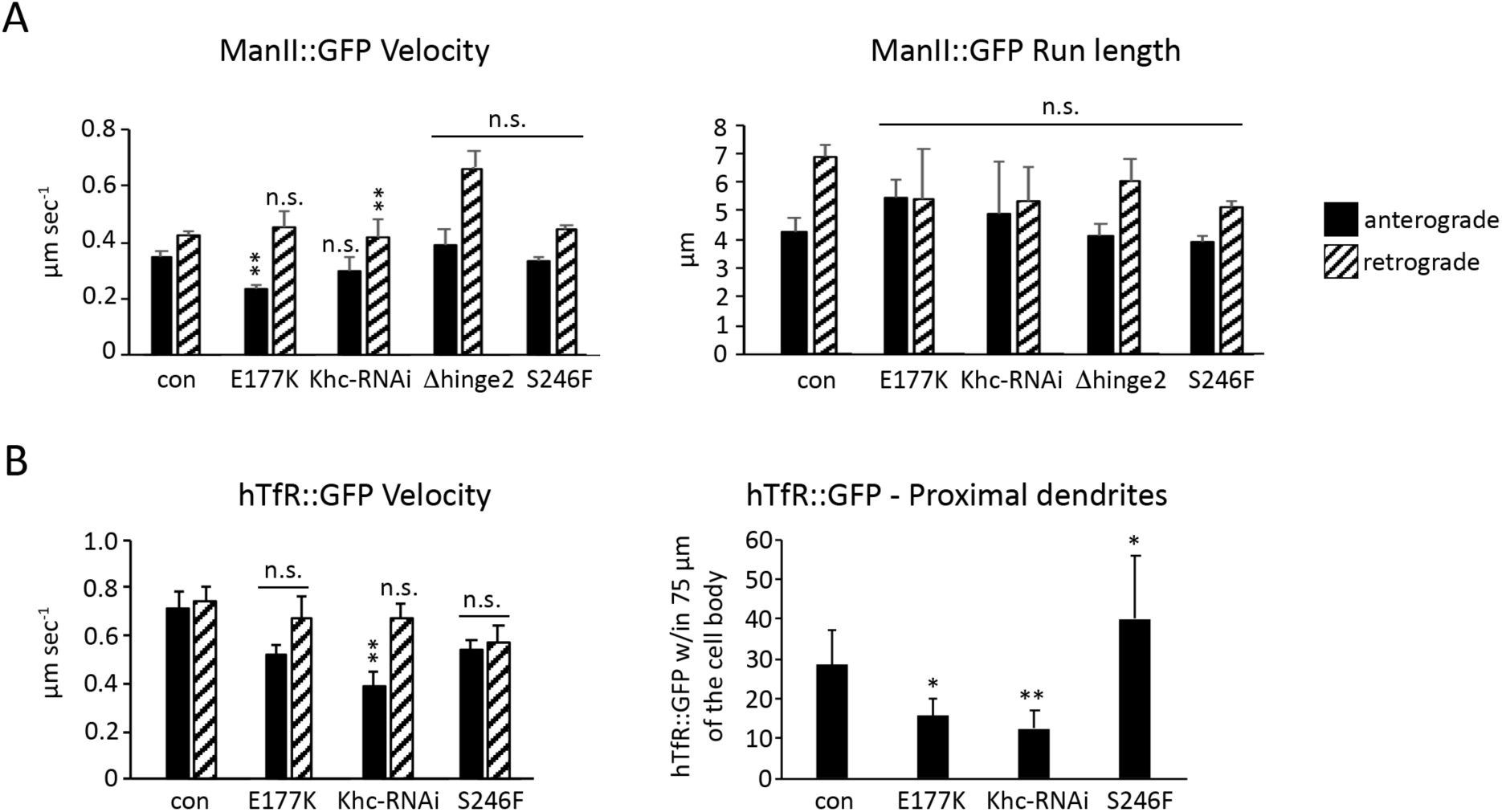
Golgi outpost and hTfR::GFP motility in control and Khc mutant neurons. (A,B) Velocity (A,B; average± SEM) and run length (A, average± SEM) of Manll::GFP (A) and hTfR::GFP(B) in control and mutant genotypes as indicated. The number of hTfR::GFP puncta are reduced in the proximal dendritic arbor. Manll::GFP: control = 338 events in 9 neurons; E177K = 107 events in 10 neurons; Khc-RNAi = 36 events in 11 neurons; hinge2 = 22 events in 9 neurons; and S246F = 519 events in 11 neurons. hTfR::GFP: control = 277 events in 11 neurons; El77K = 65 events in 11 neurons; Khc-RNAi = 88 events in 9 neurons; and S246F = 180 events in 9 neurons. ^*^P=0.05-0.01, ^**^P=0.01-0.001, n.s. = not significant in comparison to control and evaluated by one-way ANOVA and Tukey post-hoc test.

## Abbreviations

AEL: After egg laying
dlic: Dynein light intermediate chain
hTfR: Human Transferrin receptor
Khc: Kinesin heavy chain
Klc: Kinesin light chain
ppk: Mannosidase II (ManII) Pickpocket
sfGFP: Super-folder GFP
VNC: Ventral nerve cord
TIRF: Total internal reflection fluorescence

